# Synaptic strength dynamics at cortical synaptic pathways is encoded by vigilance states duration

**DOI:** 10.1101/2023.11.22.567828

**Authors:** Paul Marchal, Paul Salin, Mégane Missaire, Manon Rampon, Julien Carponcy, Régis Parmentier, Gina Poe, Gaël Malleret, Jean-Christophe Comte

## Abstract

Interactions among brain areas are essential to most cognitive functions. Neuronal interactions between these areas depend on the modulation of synaptic strength. However, this modulation remains poorly understood. We recorded evoked synaptic responses at four hippocampal pathways in freely moving male rats across 24 hours: the Perforant Path to Dentate Gyrus (*PP-DG*), Fornix to Prefrontal Cortex (*Fx-PFC*), Fornix to Nucleus Accumbens (*Fx-NAc*), and the Schaffer Collaterals to *CA1* (*SC-CA1)*. We preserved the temporal dynamics of vigilance states and synaptic responses and show for the first time that synaptic strength at these four hippocampal pathways oscillates, with a very slow periodicity. We demonstrate that synaptic strength at the *PP-DG, Fx-PFC, Fx-NAc* pathways show a positive correlation with the duration of active wakefulness (aWK) and a negative one with the duration of most sleep states (slow wave sleep (SWS) and rapid eye movement sleep (REM)), with a positive peak correlation time-lag of 1 to 10 minutes for aWK and SWS at these 3 pathways. In contrast, no significant correlation peak is found at the *SC-CA1* pathway. Finally, a model based on hypnogram data and synaptic strength at the *PP-DG* pathway was able to predict the evolution of synaptic strength at the *PP-DG, Fx-PFC* and *Fx-NAc* pathways, but not at the *SC-CA1* pathway. These results reveal that the temporal succession of vigilance states, particularly aWK and SWS, may contribute to memory processes through rapid modulation of synaptic strength at several pathways during the sleep-wakefulness cycle, suggesting that memory processes are not only dependent on sleep amount but also on sleep architecture.

## Introduction

Interactions among brain areas are essential to cognitive functions. Neuronal interactions between brain areas largely depend on synaptic functioning and its modulation. However, the modulation of synaptic strength between brain areas is not yet fully elucidated. It is thought that synaptic strength is influenced by brain states: patterns of distributed activities across the brain that depend on the animal’s physiological state. Studies have shown that vigilance states, one of the best-characterized of the brain states, are extremely dynamic. For instance, wakefulness shifts continuously between active wakefulness (aWK - with high muscle activity) and quiet wakefulness (qWK - with low EMG activity) (Crochet & Petersen, 2006; Fernandez et al., 2016; Petersen et al., 2003), while non-rapid-eye-movement (NREM) sleep alternates with wakefulness and with rapid-eye-movement (REM) sleep. Within NREM sleep, both slow wave sleep and a transitory state known as intermediate sleep (IS) occur, with IS often preceding REM sleep (Gervasoni et al., 2004). All these vigilance states are organized in cycles of several periodicities, in particular the circadian cycle but also the ultradian sleep-wakefulness cycle, showing an alternation between NREM and REM sleep within a period of 24h (Stephenson et al., 2012).

It has long been known that vigilance states modulate synaptic strength and plasticity (Bramham & Srebro, 1989; Winson & Abzug, 1977). More recent works have proposed that sleep may thus homeostatically regulate synaptic transmission through a progressive synaptic downscaling occurring particularly during SWS (a process thought to be linked to slow-wave activity) (Tononi & Cirelli, 2006). Although electrophysiological studies are in agreement with this synaptic homeostasis hypothesis (SHY), particularly those concerning interactions between neocortical areas, research on thalamo-cortical projections in the somato-sensory and visual systems seem to refute it (de Vivo et al., 2017). We have recently shown, by studying several brain connections linked to the hippocampus, that the sleep-wakefulness cycle may modulate synaptic strength at two different time scales, which could reconcile the conclusions of these different studies. At the Perforant Path to Dentate Gyrus (*PP-DG*) pathway, for example, we found that sleep reduced synaptic strength at both a sub-minute time scale, and at a slower time scale that depend on the sleep amount accumulated during the preceding 30 minutes (Rampon et al., 2023). These results suggest that vigilance states, sleep states in particular, may regulate synaptic strength in a complex manner.

These studies typically compare the average synaptic strength of each vigilance state, leaving aside the temporal dynamics between states. Notably, they do not consider the sequence of vigilance states defining the ultradian sleep-wakefulness cycle. Therefore, to tackle this issue, we performed novel analyses that preserve the temporal dynamics of vigilance states and synaptic responses and have systematically examined these dynamics in simultaneous and continuous 24h-recordings.

We thus studied the involvement of vigilance states in the modulation of synaptic strength, taking into account their relative temporal dynamics. First, we used long-term electrophysiological stimulation and recordings of field excitatory post-synaptic potentials (fEPSPs) at the *PP-DG* pathway and measured the initial slope of the synaptic response to the stimulation as a proxy for synaptic strength. We cross-correlated synaptic slopes to the average duration of each vigilance state. The correlation coefficients and time-lags were used to build prediction models. All models were extremely accurate to predict the temporal evolution of synaptic slopes of individual animals based on their vigilance. Our results show that, at the *PP-DG* pathway, only aWK strongly increases synaptic slopes, while all the other vigilance states, especially SWS and REM sleep, decreases them. Then, we focused on a panel of different pathways to assess the consistency, at the brain level, of the phenomena observed at the *PP-DG* pathway. Our results suggest that although the relationships described at the *PP-DG* pathway between synaptic strength and sleep cycle seem to be maintained at some pathways (*Fx-PFC* and *Fx-NAc*, for example), it is not the case for all four tested pathways (*SC-CA1*).

## Methods

### Animals

Data were collected from 48 Dark Agouti male rats aged of 10-15 weeks and weighing between 200-250 grams. Animals were maintained in individual cages on a 12hr/12hr light-dark cycle (9 AM-9 PM) at a room temperature of 24°C with food and water ad libitum. Light during the light phase was kept at medium intensity to minimize the interference of bright-light stress on sleep and the dark-phase rebound of REM. The animal care and treatment procedures were in accordance with the regulations of the local Lyon 1 University CE2A-UCBL 55) and European Union Council (2010/63/EU) ethics committee for the use of experimental animals. The protocol was approved by the French ethical committee for the use of experimental animals (Permit Number: DR2016-29). Efforts were made to minimize the number of animals and any pain and discomfort occurring during surgical and behavioral procedures

### Electrode Implantation

For recordings in the *Dentate Gyrus*, stereotaxic surgery was performed on 35 Dark Agouti male rats aged 10 weeks. Surgical procedures are described in detail in (Missaire et al., 2021). A recording array of 8 local field potential (LFP) electrodes was implanted in the molecular layer of the Dentate Gyrus (*DG*), and a bipolar stimulating electrode was implanted in the Perforant Path (*PP*), coming from the entorhinal cortex. In addition, 2 EEG electrodes were fixed on the skull (prefrontal and parietal), 2 EMG electrodes were inserted between the neck muscles, and a reference electrode was fixed above the cerebellum for referential recordings (**Figure 1**).

**Figure 1:**
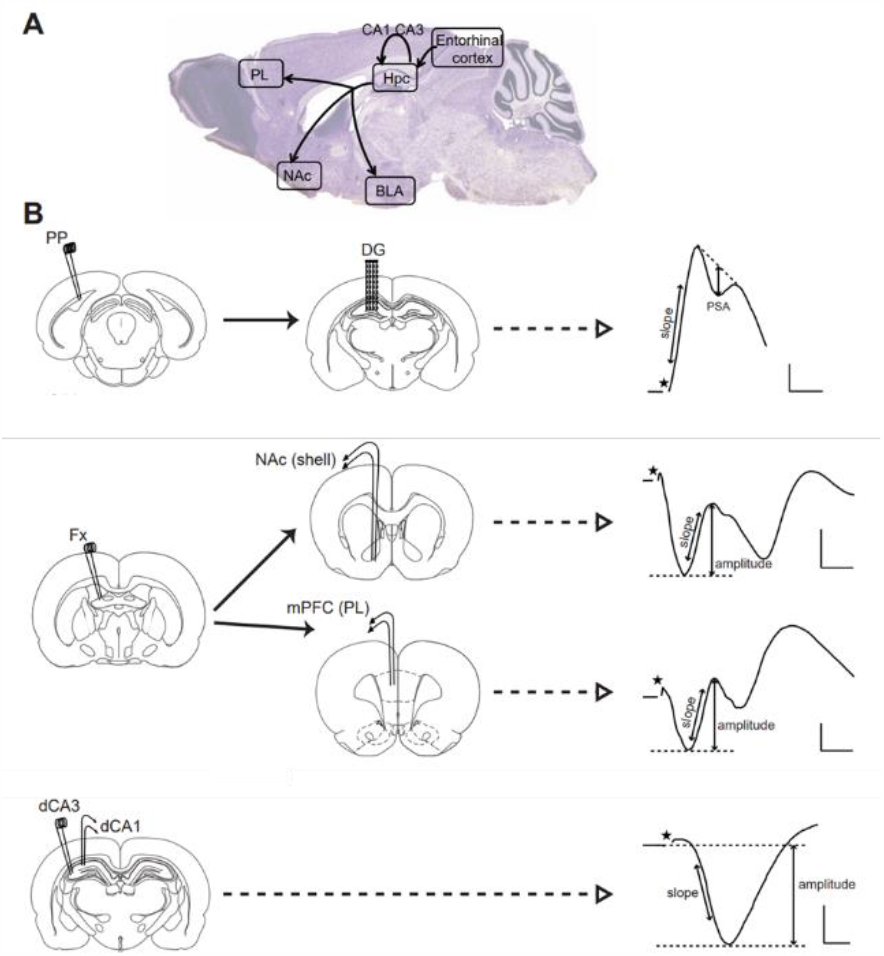
Electrode implantations and synaptic slope measurements. (**A**) Localization of the four different synaptic pathways on a sagittal view of a rat’s brain, represented by black arrows. Entorhinal Cortex to Hpc (hippocampus): PP-DG pathway. Arrow to NAc: Fx-NAc pathway. Arrow to PL: Fx-PFC pathway. CA3 to CA1: SC-CA1 pathway (**B**) For each of the studied pathways, from left to right, stimulation electrodes implantation, recording electrodes implantation localization on a transversal view of a rat’s brain, and synaptic response typical shape and slope measurement.

For recordings in the other brain regions, stereotaxic surgery was performed on 17 Dark Agouti male rats aged 10 weeks. Surgical procedures are described in detail in (Rampon et al., 2023). Four separate recording electrodes (consisting of two twisted tungsten wires) were implanted in the Prefrontal Cortex (*PFC*), Nucleus Accumbens (*NAc*) and *CA1* region of the hippocampus in each animal. Two separate stimulating electrodes were implanted in the Fornix (*Fx*) and *CA3* region of the hippocampus in each animal (**Figure 1**).

### Vigilance state classification

Vigilance states [active wakefulness (aWK), quiet wakefulness (qWK), slow wave sleep (SWS), intermediate sleep (IS – the short transient state between SWS and REM sleep) and rapid eye movement sleep (REM)] were manually scored by bouts of 5 s, based on EEG (frontoparietal differential) or LFP, and EMG recordings inspected offline, as described in (Rampon et al., 2023)(See **Figure S10** for representative examples). Briefly, aWK was characterized by a theta rhythm in CA1, DG and EEG signals, and prominent muscular activity on the EMG. qWK was characterized by weak muscular activity and delta-like activity in the PFC, NAc and LFP recordings. SWS was characterized by the occurrence of slow waves in PFC, NAc and EEG signals and a very weak muscle activity. REM sleep was determined by a prominent theta rhythm in CA1, DG and frontoparietal recordings and muscle atonia on EMG. IS was characterized by a combination of slow waves and theta rhythm in PFC and EEG recordings, along with muscle tone declining toward atonia. Recording periods that did not fulfill the criteria for any of these vigilance states were discarded. The percentage of recording time eliminated from the analyses was in average 8.2% (n=52 rats). The 24h time series (average amount of time spent in each vigilance state) was then computed over a symmetrical sliding window of 30 minutes (See **Figure 4A** for a representative example).

### Recording Setup and Electrical Stimulations

The details of the recording setups are described in (Missaire et al., 2021) and (Rampon et al., 2023). Rats remained connected to the setup during the whole duration of the experimental protocols. Electrical stimulations consisted of a monophasic 200 μs pulse delivered by an isolated pulse stimulator (model 2100, AM-Systems, USA). Before all experimental protocols, an input–output curve was established to choose the stimulation intensity used for the whole experiment. This intensity had to elicit a fEPSP with an amplitude corresponding to 75% of the maximal amplitude on the electrodes. During the experimental protocol, stimulations were elicited every 30s for 24h. Signal was acquired at a frequency of 5000Hz.

### Synaptic responses and slope measurement

The triggered synaptic responses at the different pathways were extracted from the raw signal recorded by each electrode (**Figure 2**). The initial part of the slope (a window of 2 ms from the first minimum of the response) of each synaptic response was fitted by a 3^rd^ order polynomial.

**Figure 2:**
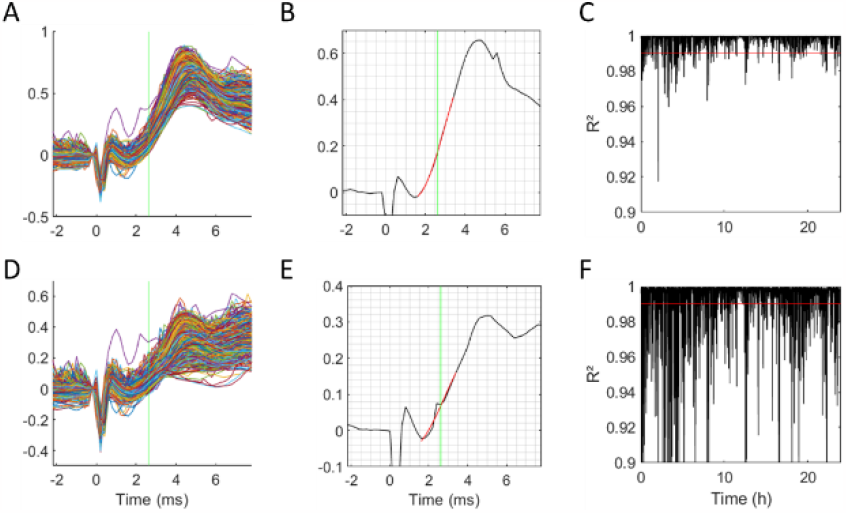
Synaptic slopes, first derivative of the 3^rd^ order polynomial fit. (**A, D**) Examples of stacked synaptic responses measured at two different electrodes in the Dentate Gyrus, during 24h. The signal is normalized at t=0 (stimulation). The synaptic response slopes are measured in the initial portion of the response (at 2.5 ms, green vertical line). (**B, E**) Examples of 3^rd^ order polynomial fits of two synaptic responses from electrodes shown in A and D, respectively. Synaptic responses and polynomial fits are respectively represented in black and red, while the green vertical line represents the time point at which synaptic response slope is calculated. (**C, F**) R^2^ coefficients for each of the polynomial fits of the synaptic responses presented in A and D, respectively. The red horizontal line represents the exclusion threshold (R^2^=0.99) below which individual measures were excluded. (**A-C**) Examples of valid recordings (with <10% of unreliable slopes). (**D-F**) Examples of excluded recordings or electrodes. (**E**) Example of an excluded synaptic response slope measure (with R^2^>0.99). (**F**) Example of an excluded electrode (with >10% of unreliable slopes).

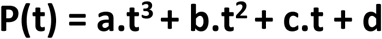

The synaptic response slope (value of the first derivative of the polynomial regression) was analytically calculated at t = 1ms (**Figure 2B**).

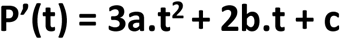

A fit quality measurement (R^2^) was performed for each synaptic response. Polynomial fits leading to unreliable slope measures (R^2^<0.99) were systematically excluded (**Figure 2E**). This method allows a more reliable measure of synaptic slopes, as suggested by several randomized tests on different pathways. By applying this method, it is possible to reliably remove artefactual measurements (see **Figure 2**).

For measures of fEPSP slope at the PP-DG pathway, recordings with more than 10% of unreliable slopes were excluded (**Figure 2F**). For each rat, from 2 to 8 electrodes were used to calculate the average synaptic slope of each synaptic response. The 24h time series was then smoothed by a symmetrical rectangular sliding window of 30min width (see **Figure 4A** for a representative example).

For measures of fEPSP slope at the *Fx-PFC, Fx-NAc* and *SC-CA1* pathways, the decision of including each recording site was taken based on histological inspection of the electrodes’ positions as well as the shape and stability of synaptic responses, as described in (Rampon et al., 2023). The 24h time series was then smoothed by a symmetrical rectangular sliding window of 30min width (see **Figure S17A**, **Figure S18A**, **Figure S22A** for representative examples).

### Signal pre-processing

Each time series (synaptic slopes or vigilance states) was z-scored (subtracted of its mean and divided by its standard deviation). Additionally, the trend of each time series was removed (subtraction of the best fitted 3^rd^ degree polynomial) to avoid correlations or cross-correlations artefacts in subsequent analyses (see **Figure 4**B for a representative example).

### Prediction model

We constructed a synaptic response slope dynamics prediction model, using the Wiener filter procedure (Noble & Weiss, 1959). To construct the following model, cross-correlations were performed between synaptic response slope signals recorded over 24h and each vigilance state average durations (aWK, qWK, SWS, IS, REM) (see **Figure 5**C for representative examples). Optimal (maximal or minimal) cross-correlation coefficients (C_x_) and their corresponding time-lag (t_x_) were used to build synaptic response slope (S) prediction models (**Figure 3**).

**Figure 3:**
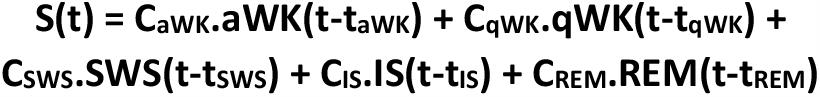
Predictive model equation. Models were designed to predict synaptic response slopes dynamics (S(t)) based on vigilance states dynamics (aWK(t), qWK(t), SWS(t), IS(t) and REM(t)). Each dynamic element of the models was weighted by a corresponding cross-correlation coefficient (C_x_) and delayed by a corresponding time-lag (t_x_).

**Figure 4:**
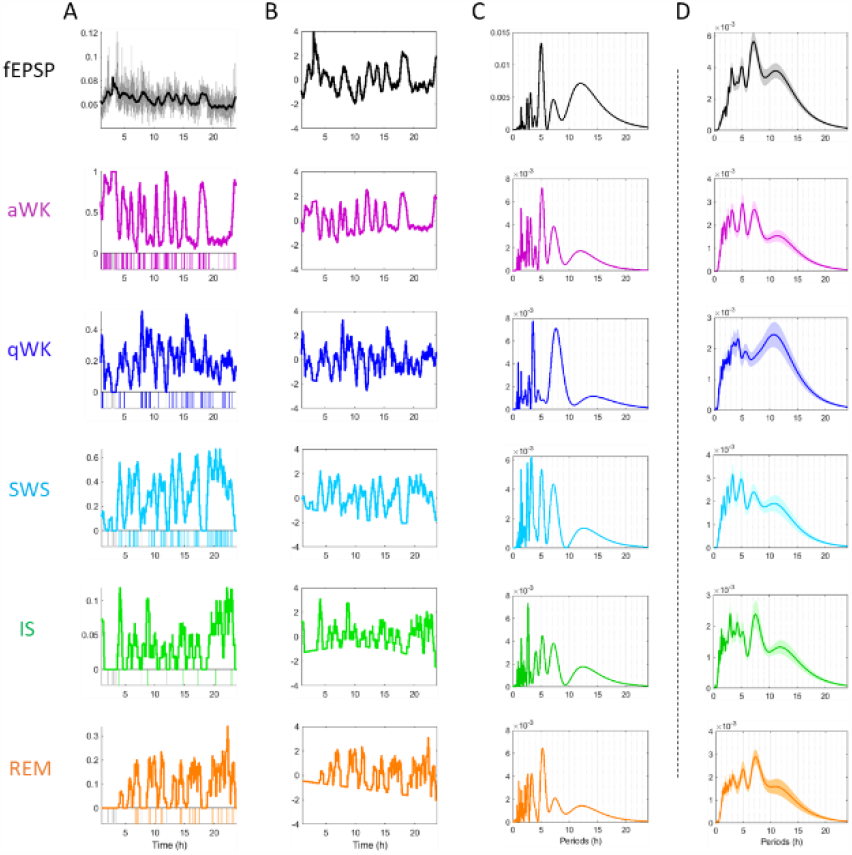
Synaptic response slopes and vigilance states share common rhythms. (**A**), (**B**) From top to bottom: synaptic response slopes at the PP-DG pathway (fEPSP, in black) and vigilance states occurrence (aWK = active Wakefulness, in purple; qWK = Quiet Wakefulness, in dark blue; SWS = Slow Wave Sleep, in light blue; IS = Intermediate Sleep, in green; REM = Rapid Eye Movement Sleep, in orange) for a representative animal for 24 hours. (A) For synaptic slopes, the grey line represents raw measures, and the black line represents its sliding average over 1h. For vigilance states, the bottom histograms represent bouts of each state in color and missing values in grey. The colored lines represent the sliding averages of each vigilance state over 1h. (**B**) Black and colored lines represent normalized and detrended sliding averages. (**C**) Periodograms of the signals in B. (**D**) Mean periodograms for all rats (n=35). Shaded areas represent SEM.

**Figure 5:**
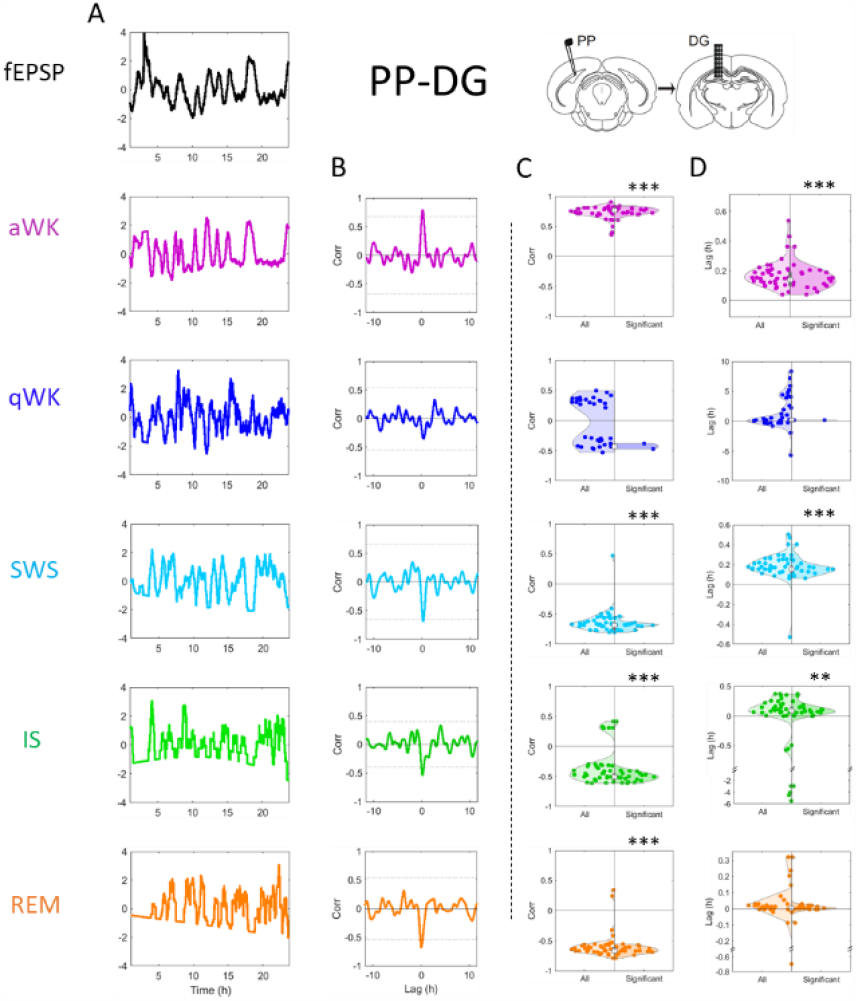
Vigilance states predict synaptic strength dynamics at the PP-DG pathway. (Color code like figure 4). (**A**) The black and colored curves represent the normalized and detrended synaptic response slopes and average duration of vigilance states signals, respectively. (A) Cross-correlations between synaptic response slopes and the average duration of each vigilance state for a representative animal. The horizontal dotted lines represent the significance threshold for each cross-correlation. (**C**) Cross-correlation coefficients between synaptic response slopes and vigilance states for all rats (n=35). (**D**) Time-lags in the cross-correlations between synaptic response slopes and vigilance states for all rats (n=35). (**C-D**) Right side of each panel shows only the significant cross-correlation coefficients and their corresponding time-lags, without spurious correlations.

The model was built to determine whether different cortical pathways shared a similar relation between vigilance states and synaptic strength. Thus, the model was built based on the median parameters (correlation factors and related time-lags) of data measured at the *PP-DG* pathway. The model fidelity was assessed by calculating the correlation coefficient between the measured synaptic slopes and the model-predicted synaptic slopes (see **Figure 9** for a representative example).

### Statistical analysis

Signal processing and statistical analyses were performed using Matlab™ home-made scripts. Cross-correlation significance limits (ccsl) were calculated with the following expression, where a and b represent the autocorrelation coefficients of the 2 time-series considered for cross-correlation, respectively, and n represents the number of samples of the time series. This expression corrects the traditional ccsl formula to avoid spurious correlations due to autocorrelation (Dean & Dunsmuir, 2016).

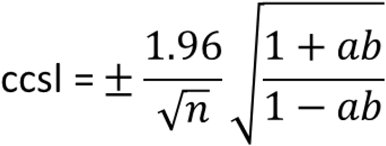

Statistical differences between correlation coefficient or time-lag and the theoretical mean of 0 or between correlation coefficients were assessed using one-sample t test (or Wilcoxon signed rank test if normality criterion was not met). Statistical differences between time-lags were assessed using paired t-tests (for these tests, only animals with significant time-lags for all vigilance states were used, hence the reduced sample size).

## Results

### Slow periodic modulation of synaptic strength at the PP-DG pathway

The dynamics of synaptic strength at the PP-DG pathway were quantified as synaptic slopes of fEPSP on 24h continuous recordings. As shown on a typical rat, slow periodicities were observed (**Figure 4A,B**). In this quantification, given that synaptic responses are highly variable in slope, in order to avoid the effect of large variations in synaptic strength, we used a sliding average over 30 minutes. Periodograms (**Figure 4C**) revealed that synaptic strength mainly oscillated with a single period. On the population of 35 rats as a whole, an average peak was observed at 7h, as well as other less prominent peaks at 3, 5 and 11 hours (**Figure 5D**).

The sleep-wake cycle exhibits an ultradian rhythm (Stephenson et al., 2012, 2013). We thus hypothesized that a relationship between ultradian sleep-wake cycle and synaptic strength modulations might also exist. Therefore, we represented the dynamics of vigilance states using the same sliding average method as for synaptic strength. Sliding averages of binary coded vigilance states allowed a computation of the average amount of time spent in each vigilance state (**Figure S11**). In this quantification of vigilance states, we have separated wakefulness into quiet wakefulness (qWK) and active wakefulness (aWK), and considered the transition between SWS and REM sleep, intermediate sleep (IS), to be a sleep state different from the other two (see Methods). We showed the periodicity of the vigilance states in the same rat (**Figure 4A,B**). Periodograms revealed that aWK, SWS and REM oscillate with periods of approximately 3, 5 and 11 hours (**Figure 4C,D**). These periodicities of vigilance durations are close to those of synaptic strength. However, as shown in **Figure 4D**, the periodicity was different for qWK compared to the other vigilance states.

Based on these results, we next hypothesized that a temporal relationship might exist between synaptic strength and vigilance states dynamics. To test this hypothesis, we cross-correlated the amounts of each vigilance states and the strength of the synaptic response for each rat (**Figure 5**).

We found that vigilance states were correlated to the synaptic strength measured at the *PP-DG* pathway (**Figure 5B,C**). Most correlations ranged from strong (>0.75) to moderate (0.5-0.75) and were positive for aWK but negative for the three sleep states (**Figure 5C**). Given that the modulation of the amounts of vigilance and synaptic strength were both periodic over 24 hours, there was a risk of obtaining artifactual peaks of correlation. For this reason, we computed surrogates for the analyses obtained for each vigilance state and for each animal. By computing surrogates (thin horizontal lines shown in **Figure 5B**, see details in Methods) we observe that most of these correlations were significant (see the right-hand side of the graphs in **Figure 5C**, **Table S1**). Interestingly, the correlation peaks showed a positive time-lag of several minutes with respect to time 0 for aWK and SWS (aWK: 8.7 min, p<0.001; SWS: 10 min, p<0.001, **Table S2**). This means that an increase in aWK duration precedes an increase in synaptic strength, while an increase in SWS duration precedes a decrease in synaptic strength. An analysis of the time-lag differences for each animal showed that SWS amount precedes a decrease in synaptic strength by a significantly longer time than aWK amount precedes an increase in synaptic strength (aWK vs SWS: p<0.05, n=14, **Figure S23A**). For REM, on the other hand, the time-lag is on average close to time 0 (**Figure 5D**, **Table S2**). Thus, a modulation of REM amount seems almost synchronous to the modulation of synaptic strength.

In contrast with the other vigilance states, qWK amount, did not correlate with synaptic strength (only 2 out of 35 animals correlated significantly, **Table S1** and **Table S2**). Thus, a modulation of qWK amount does not seem to follow or precede a change in synaptic strength. Taken together, these results suggest that a change in synaptic strength at the *PP-DG* pathway can be predicted several minutes in advance by quantifying the amount of SWS and aWK, but not by quantifying that of qWK and REM.

Studying synaptic strength modulation at the *PP-DG* pathway revealed a strong temporal link between the ultradian sleep-wakefulness cycle and synaptic strength at this pathway. In the upcoming sections, we will see that this conclusion may apply to some other cortical pathways linked to the hippocampus, but not all.

### Correlations at the Fornix to Prefrontal Cortex pathway

We analyzed the modulation of synaptic strength at the Fornix to Prefrontal Cortex pathway (*Fx-PFC*, connections mainly linking the hippocampus to the medial Prefrontal Cortex, **Figure 6**). We observed, as in the case of the *PP-DG* pathway, a periodic variation of synaptic responses over 24h (periodicity of approx. 4h, **Figure S16**). We therefore correlated vigilance state durations with synaptic strength. We showed that the synaptic strength correlated positively with aWK and negatively with SWS and REM (**Figure 6C**, **Table S3**). The peaks of cross-correlations of aWK and SWS amount with synaptic strength had positive time-lag, but closer to time 0 than for the *PP-DG* pathway (aWK: 0.80 min, p=0.15; SWS: 2.3 min, p<0.05, **Table S4**). However, peaks of cross-correlations of REM amount with synaptic strength showed negative time-lag (-7.8 min; p<0.001, **Table S4**). As for the *PP-DG* pathway, qWK amount did not seem to correlate (only 2 out of 9 animals correlated significantly) with synaptic strength (**Table S3, Table S4**). Surprisingly, IS did not correlate with synaptic strength (**Table S3**). These results suggest that the modulation of aWK, SWS and REM durations predicts the dynamic modulation of synaptic strength at the *Fx-PFC* pathway, in a similar fashion as the one observed at the *PP-DG* pathway.

**Figure 6:**
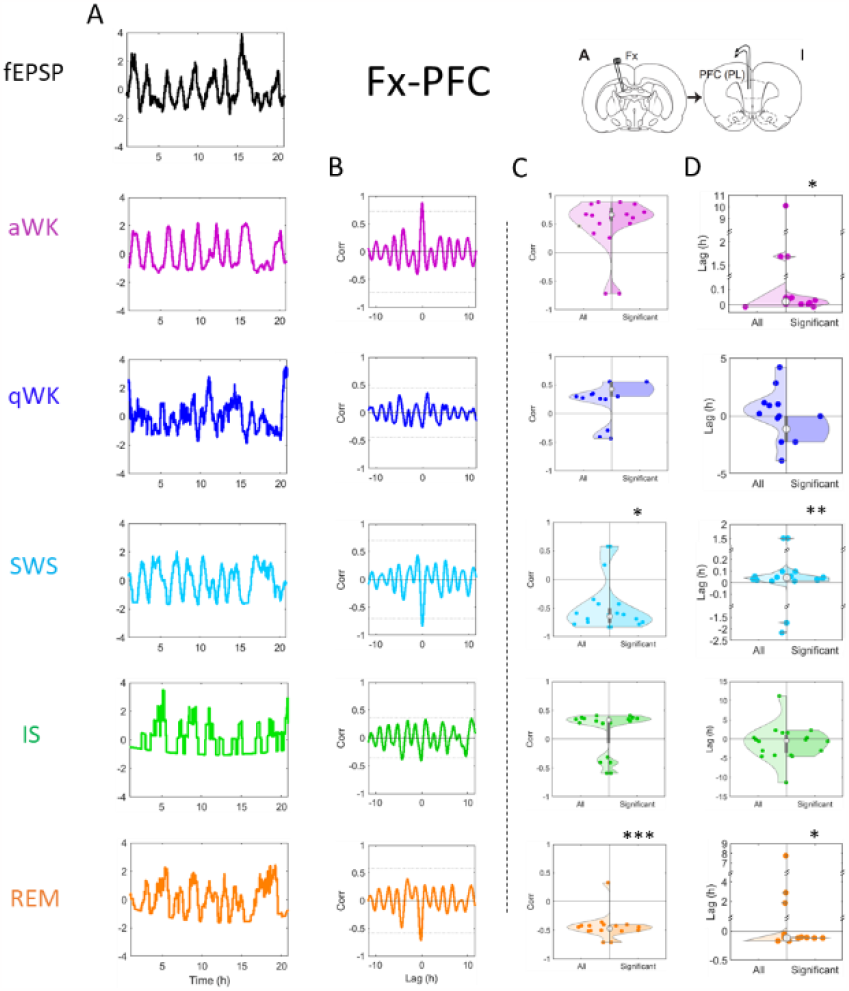
Vigilance states predict synaptic strength dynamics at the Fx-PFC pathway. See Figure 5 for legend (n=10).

### Correlations at the Fornix to Nucleus Accumbens pathway

At the Fornix to Nucleus Accumbens pathway (*Fx-NAc*, **Figure 7**), we also observed a slow periodic variation of synaptic strength over 24h (periodicity of approx. 5h, **Figure S17**). In addition, we showed that aWK, SWS and REM amount correlated with synaptic strength (**Figure 7C**). Indeed, aWK tends to positively (and in most cases, 7 out of 9 animals, significantly) correlate with synaptic slopes and SWS and REM tend to negatively correlate with synaptic slopes (**Table S5**). The peaks of cross-correlations for aWK or and SWS were close to 0 time-lag (aWK: 0.36 min, p=0.77; SWS: 0.40 min, p=0.79, **Table S6**). However, like at the *Fx-PFC* pathway, the peak of cross-correlations between synaptic slopes and REM showed a negative time-lag (-6.9 min, p<0.001, **Table S6**). qWK amount did not seem to correlate (none of the 9 animals correlated significantly) with synaptic strength (**Table S6**). These results suggest that the dynamic modulation of aWK, SWS and REM durations is temporally linked with the dynamic modulation of synaptic strength at the *Fx-NAc* pathway.

**Figure 7:**
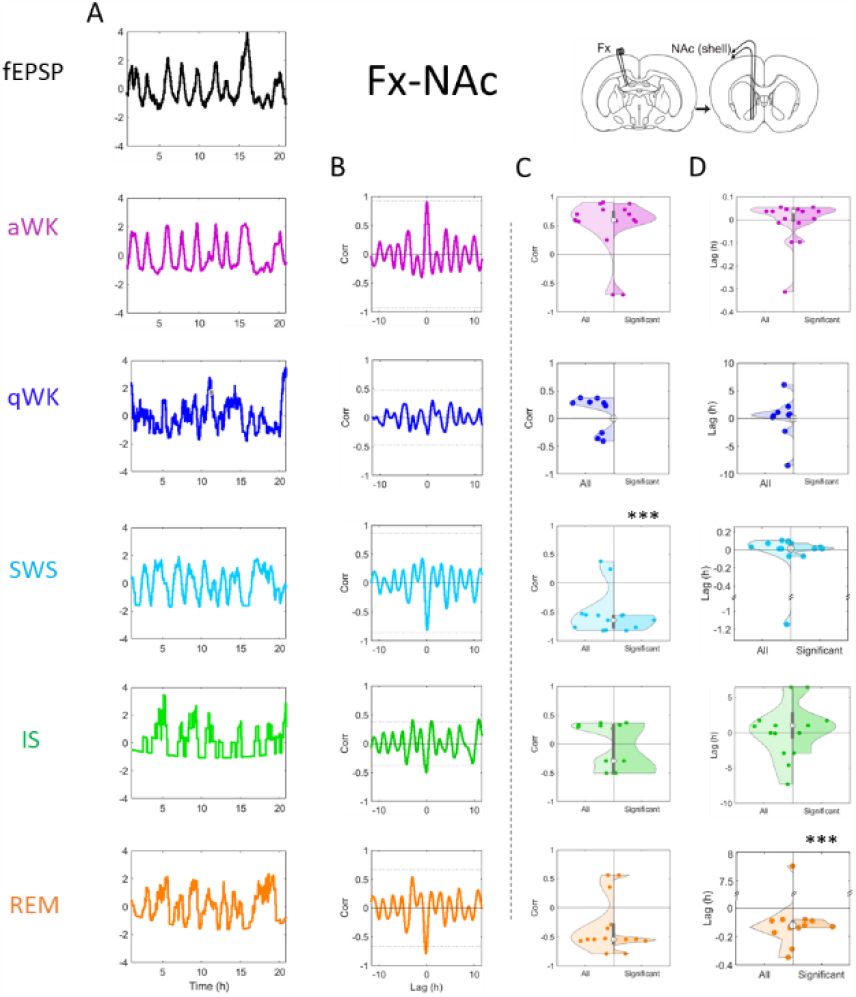
Vigilance states predict synaptic strength dynamics at the Fx-NAc pathway. See Figure 5 for legend (n=9).

### Absence of correlation at the Schaffer Collaterals to CA1 pathway

Finally, at the Schaffer Collaterals to CA1 pathway (*SC-CA1*, **Figure 8**), we found that, unlike previous pathways, synaptic strength variations were dominated by very slow oscillations of around 12 hours (**Figure S18**). However, also in contrast with the other pathways, no significant peak of cross-correlation was observed with the different vigilance states (**Figure S21, Table S7**). These results suggest that vigilance state amount does not follow or precede any kind of modulation of synaptic strength at this pathway.

**Figure 8:**
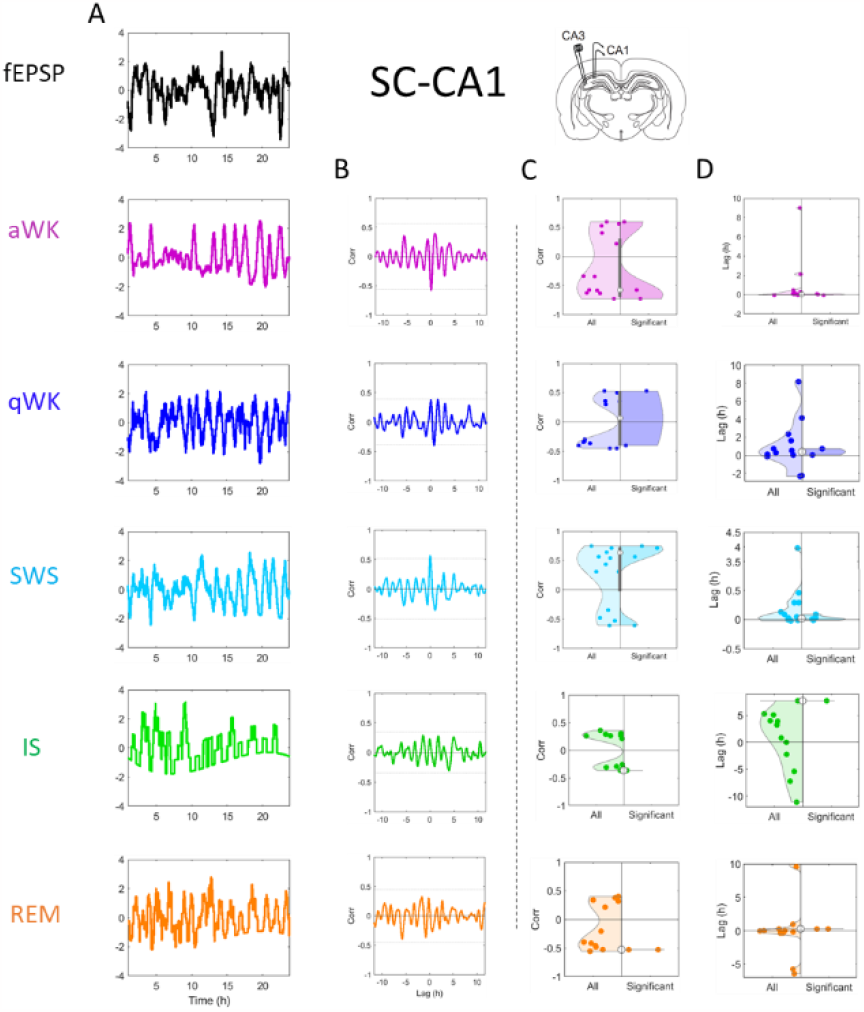
Vigilance states do not predict synaptic strength dynamics at the SC-CA1 pathway. See Figure 5 for legend (n=12).

Altogether, our results show that a relationship seems to exist between the sleep-wakefulness cycle and the strength of the synaptic response at three out of the four pathways we studied. We next hypothesized that this relationship might be similar at these three pathways, which would suggest that a common mechanism might be responsible for these relationships across multiple pathways of the brain. To test this hypothesis, we built a computational model of the relationship between synaptic strength and vigilance states.

### Prediction model

Since a strong correlation exists at some pathways between the amount of vigilance states and the strength of the synaptic response, we built a model aiming at predicting the synaptic response variations based on the sleep-wake cycle. We used the median values of the significant correlation coefficients and the corresponding time-lags of cross-correlations between synaptic response slopes and vigilance states (aWK, qWK, SWS, IS and REM) at the *PP-DG* pathway (see Methods **Figure 3A**) to predict synaptic responses at the *PP-DG* pathway for each rat (see **Figure 9A** for a representative example). We then correlated measured and predicted values of synaptic strength to assess the predictivity of the model (see **Figure 9B** for a representative example).

**Figure 9:**
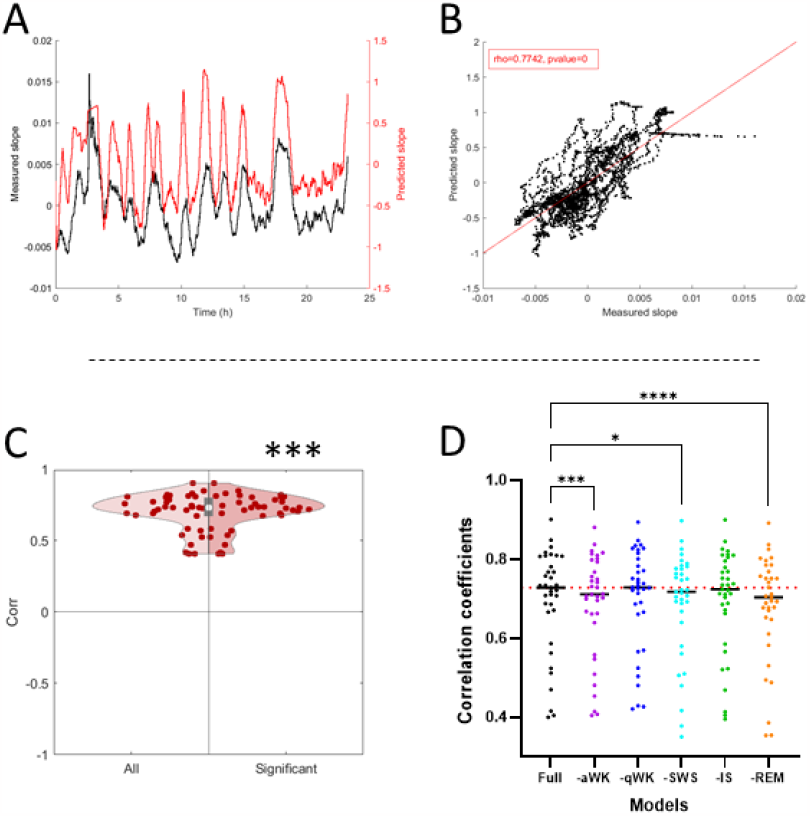
Vigilance state durations predict synaptic strength dynamics at the PP-DG pathway. (**A**) Slope prediction based on vigilance states for a representative animal. The black curve represents synaptic response slopes measured at the PP-DG pathway. The red curve represents synaptic response slopes predicted by the model. (**B**) Correlations between measured slopes and predicted slopes. The red line represents the least-square line for the correlation. (**C**) Correlation coefficients between measured slopes and predicted slopes for all rats (n=35). Right side of the panel shows only the significant correlation coefficients. Difference to the theoretical mean of 0 was assessed by one sample t-test. ***: p<0.001. (D) Correlation coefficients between measured slopes and predicted slopes by each model for all rats (n=35). Black dots represent individual correlation coefficients with the full model (including all vigilance states). Purple dots with the partial model including all vigilance states except aWK. Dark blue dots with the partial model including all vigilance states except qWK. Light blue dots with the partial model including all vigilance states except SWS. Green dots with the partial model including all vigilance states except IS. Orange dots with the partial model including all vigilance states except REM. Difference between models was assessed by Friedman’s test. Differences of each model to the full model were assessed by Dunn’s test. *: p<0.05. **: p<0.01. ***: p<0.001.

Correlations between measured and predicted synaptic response slopes revealed that the model explains around 70% of the variations in synaptic response slopes at the *PP-DG* pathway of our population of rats (**Figure 9C**, n=35, C=0.73, p<0.001). This result confirmed that vigilance state amounts can predict synaptic strength variations at the *PP-DG* pathway.

However, we previously showed that qWK does not significantly correlate with synaptic response slopes at the *PP-DG* pathway (**Figure 5**). We thus wondered if including this vigilance state in the model would in fact decrease its predictive capability. To test this assumption, we decided to build 5 additional models, each omitting only one vigilance state. We then assessed the predictivity of these models and compared it to that of the full model (**Figure 9D**).

This analysis revealed that removing either aWK, SWS or REM from the full model significantly decreased its predictivity. However, removing either qWK or IS did not. These results suggest that while aWK, SWS and REM each play a crucial role in the modulation of synaptic strength, neither qWK nor IS do.

We then assessed if the full model, built based on data measured at the *PP-DG* pathway, could accurately predict synaptic response slopes at other pathways. We used the same model to predict synaptic response slopes at the *FX-PFC, Fx-NAc* and *SC-CA1* pathways (see **Figure S19A**, **Figure S20A**, **Figure S21A** for representative examples respectively) and correlated measured and predicted values for synaptic response slopes to assess the predictivity of the model at each pathway (see **Figure S19B**, **Figure S20B**, **Figure S21B** for representative examples).

We show that the full *PP-DG* model explains significant amounts (50% and 40%) of synaptic response slopes variations at the *Fx-PFC* and *Fx-NAc* pathways, respectively (**Figure S19C**: n=9, C=0.54, p<0.001; **Figure S20C**: n=9, C=0.40, p<0.05). However, it does not seem to explain significant amounts of synaptic response slopes variations at the *SC-CA1* pathway (**Figure S21C**: n=12, C=0.33, p=0.27). Taken together, these results suggest that the dynamics of the sleep-wakefulness cycle can explain the modulation of the synaptic strength at the *PP-DG, Fx-PFC and Fx-NAc* pathways but not at the SC-CA1 pathway.

## Discussion

In this study, we demonstrate for the first time, using a sliding average method, that synaptic strength oscillates with a very slow periodicity of 4-7h at four hippocampal pathways. Since the average duration of vigilance states also tends to oscillate at similar periodicities, we analyzed the temporal link potentially existing between these and the strength of the synaptic responses. Our results show that, at the Perforant Path to Dentate Gyrus (*PP-DG*), Fornix to Prefrontal Cortex (*Fx-PFC*) and Fornix to Nucleus Accumbens (*Fx-NAc*) pathways, a positive correlation exists for active wakefulness (aWK) and a negative one for most sleep states [slow wave sleep (SWS), intermediate sleep (IS) and rapid eye movement sleep (REM)]. Additionally, we found a positive peak correlation time-lag of 1 to 10 minutes for aWK and SWS at these 3 pathways. In contrast, we found no significant correlation peak at the Schaffer Collaterals to *CA1* (*SC-CA1*) pathway. Finally, a model based on the analysis of sleep-wakefulness cycle data indicates that the amounts of wakefulness and sleep are able to predict the evolution of synaptic strength at the *PP-DG, Fx-PFC* and *Fx-NAc* pathways, but not at the *SC-CA1* pathway. Below we first discuss the methods used in the present work and examine the relevance of these findings for understanding the role of synaptic changes in cognitive functions.

In the present work we have developed several analysis tools, including one that substantially improves the quantification of the slope of fEPSPs. We introduced a novel procedure to accurately and reproducibly measure synaptic response slopes, using polynomial fits. Indeed, this method allows for automatic exclusion of unreliable measures, thus improving the signal/noise ratio. We then used a sliding average calculated over a time window of 30 min. We chose a sufficiently large window to calculate a meaningful average, and sufficiently short to do not smooth dynamic phenomena compatible with the dynamics of vigilance states. The sliding average method allows for transforming binomial data (vigilance states scoring) into continuous data (average vigilance states duration, **Figure S11**) and thus allows for a temporal dynamic preserving analysis (cross-correlations). The fact that we found significant time-lags in the cross correlations of vigilance states with synaptic strength illustrates the importance of using cross-correlations. Indeed, simple correlations, not controlling for time-lag, might have resulted in underestimated/erroneous correlations.

The average durations of aWK, SWS, IS and REM all share a very similar periodogram with that of synaptic response slopes, especially strong oscillation at periods of around 5-7 hours. Surprisingly, the average duration of quiet Wakefulness (qWK) shows a different pattern of oscillations to that of the other vigilance states. This implies that qWK is somehow relatively independent of the usual ultradian sleep wake cycle (aWK ➔ SWS ➔ IS ➔ REM ➔ aWK). Indeed, qWK which is a heterogeneous state that can occur almost anywhere during this cycle (qWK can be either observed after a period of strong activity (aWK) or during a period of rest, as a short awakening between two SWS episodes), explaining its particular periodogram profile.

We performed cross-correlations between synaptic response slopes and each vigilance state average duration to show that, at the *PP-DG, Fx-PFC* and *Fx-NAc* pathways, the average durations of most vigilance states are strongly correlated with synaptic strength. On one hand, we showed that the average duration of aWK positively correlates with, and often precedes, the synaptic response variations, suggesting that aWK dynamically upregulates synaptic strength. On the other hand, we showed that SWS negatively correlates with, and often precedes, the synaptic response, suggesting that this vigilance state dynamically downregulates synaptic strength. These results are reminiscent of previous demonstrations that wakefulness promotes synaptic potentiation while sleep supports synaptic depression (Bramham & Srebro, 1989; de Vivo et al., 2017; Gervasoni et al., 2004; Rampon et al., 2023). However, we additionally demonstrated here that synaptic strength modulation appears to be linked to the average duration of each vigilance state, emphasizing the importance of temporal patterns in this relationship, that is the sleep architecture.

Additionally, we showed that REM negatively correlates with the synaptic response, much like SWS does, but often with an unexpected negative time-lag. This result should logically imply that it is the synaptic slopes variations that regulate REM sleep occurrence, and not the other way around, as it is the case for SWS. However, we believe it is unlikely that synaptic strength would drive REM variations. More likely, we believe that the functional relationship between SWS and REM (REM bouts are mainly preceded by SWS bouts (Benington & Heller, 1994; Bjorness et al., 2018; Kishi et al., 2011)) could explain the apparent correlation between REM and synaptic response, as well as the negative time-lag of REM compared to SWS. Indeed, SWS and REM are strongly positively correlated, SWS largely preceding REM (**Figure S24**). At least part of the correlation observed between REM and synaptic slopes would thus be due to the strong correlations between REM and SWS and between SWS and synaptic slopes. This does not exclude however that REM can have a short-term effect on synaptic slopes, at the very transition towards this state, as demonstrated previously (Rampon et al., 2023, **Figure S22**). However, short-term effects are unlikely to weigh in our correlations because of the use of sliding averages.

It is interesting to note that the use of cross-correlations, as opposed to simple correlations, allows to not underestimate the correlations that exist between synaptic strength and vigilance states or between vigilance states. Additionally, by conserving the temporal organization of the time series, cross-correlations allow to hint at causation where simple correlations do not. On the other hand, performing sliding averages then cross correlations tends to hide the shorter-term effects (*e*.*g*., see pre above).

One of the most intriguing results of the present work is that, in contrast to aWK, the amounts of quiet wakefulness (qWK), although changing periodically, do not correlate with synaptic strength for the 4 pathways studied. qWK is a state of wakefulness well characterized in rodents by low muscle activity on EMG and slow activity close to delta oscillations in several cortical areas (such as in the PFC). In addition, separating aWK and qWK leads to clear differences in thalamic activities, as well as cognitive (such as sensory discrimination capacity) and physiological parameters (pupillary diameter). This is not to say, however, that qWK has no effect on synaptic strength, as for example, at the *PP-DG* and *SC-CA1* pathways the switch from aWK to qWK leads to sub-minute changes in synaptic responses (Rampon et al., 2023). Compared to aWK, during qWK, acetylcholine and norepinephrine signaling is less activated (Kodama et al., 2002). This weaker signaling during qWK may reduce the variability in synaptic transmission (Rampon et al., 2023) and additionally prevent the cumulative effect we observed here for aWK. During this state, hippocampal ripples can often be observed, playing an important role in replays of precise neuronal sequences and in the consolidation of spatial memory (Girardeau & Zugaro, 2011). Future studies have to determine whether qWK is a privileged period for more stable synaptic transmission (*i*.*e*., no cumulative effect) in contrast to aWK and sleep stages.

In connection with this finding, we reveal that the average duration of qWK shows a different pattern of oscillations than the one observed for the other vigilance states. This implies that qWK is somehow relatively independent of the ultradian cycle. Indeed, the cross-correlation of all vigilance states amounts (see supplementary **Figure S24**) reveals that the level of correlation is lower between qWK and the other vigilance states on one hand than between these other vigilance states on the other hand. Interestingly, by removing the qWK parameters, the degree of predictability of synaptic strength is not altered compared to what happens when the aWK, SWS and REM parameters are individually withdrawn (**Figure 9D**). Although qWK has sometimes been considered a state of falling asleep or transition between wakefulness and sleep, it is now considered a sub-state of wakefulness with its own characteristics that distinguish it from aWK and SWS. It is a brain state characterized behaviorally by a high degree of attention, during which ripple activity coupled with hippocampal activity replays can be observed. The reason for the low level of correlation we observed here between qWK and the other vigilance states is currently unclear. Studies suggest multiple states of wakefulness, with local changes in brain states depending on internal and contextual demands. In contrast, the ultradian alternation between wakefulness and SWS or REM depends on sleep pressure (as indicated by the progressive increase in slow wave activity during SWS) and competition/facilitation between SWS and REM (Benington & Heller, 1994; Bjorness et al., 2018). This might explain the low level of correlations between qWK and synaptic strength - given that aWK positively correlates to synaptic strength and sleep negatively does. Additionally, a bout of qWK occurring after a bout of aWK might not be functionally equivalent to one occurring after a bout of REM sleep (Jarosiewicz et al., 2002). This possible heterogeneity of qWK might explain the difficulty to demonstrate clear trends in correlations between qWK and synaptic strength.

At the *SC-CA1* pathway, we could not evidence clear correlations between the average duration of any vigilance state and synaptic strength like it has been observed at the three other pathways (*PP-DG, Fx-PFC* and *Fx-NAc*). These results suggest that the sleep cycle might not play as strong a role in the modulation of synaptic strength at the *SC-CA1* pathway, contrary to the one implied at the three other pathways (according to our results).

Our model accurately predicted the temporal dynamics of synaptic response slopes at the *PP-DG* pathway, based on the hypnogram of each rat. This result suggests that a same mechanism rules the relationship between the ultradian sleep wake cycle and the synaptic strength in the whole population. As we could expect, the model is based on the correlation coefficients and time-lags measured at the *PP-DG* pathway and it could very accurately predict the synaptic response slopes at this same pathway. However, the model was also able to accurately predict the synaptic response slopes variations at two other pathways (*Fx-PFC* and *Fx-NAc*, but not *SC-CA1*). These results suggests that a common mechanism, shared by all individuals, rules the relationship between the ultradian sleep wake cycle and the dynamic modulation of synaptic strength at multiple pathways across the brain, with some exceptions.

To summarize, we show in this study that the ultradian sleep cycle, particularly the alternance of aWK and SWS, might be responsible for the dynamic modulation of the synaptic strength at different pathways across the brain. We hypothesize that the broad periodic modulations of synaptic strength might create dynamic time windows during which a synaptic connection is more susceptible to changes. Highs and lows in global synaptic strength might help fine-tuning smaller scale synaptic strength. The connection between two neurons might benefit from a general downscaling at the global pathway they are a part of. This phenomenon might be crucial for synchronizing wakefulness-related experiences and learning and memory processes during sleep.

Finally, we propose that the predictive model presented in this study could be used in the future to formulate working hypotheses by forecasting theoretical changes in synaptic strength induced by artificial manipulation of vigilance states and brain oscillations. These hypotheses could then help design targeted experimental protocols to challenge such theoretical changes and thus offers a tool for studying the role of synaptic changes in different limbic networks in cognitive functions such as memory and emotion regulation.

## Supplementary Materials

**Figure S10:**
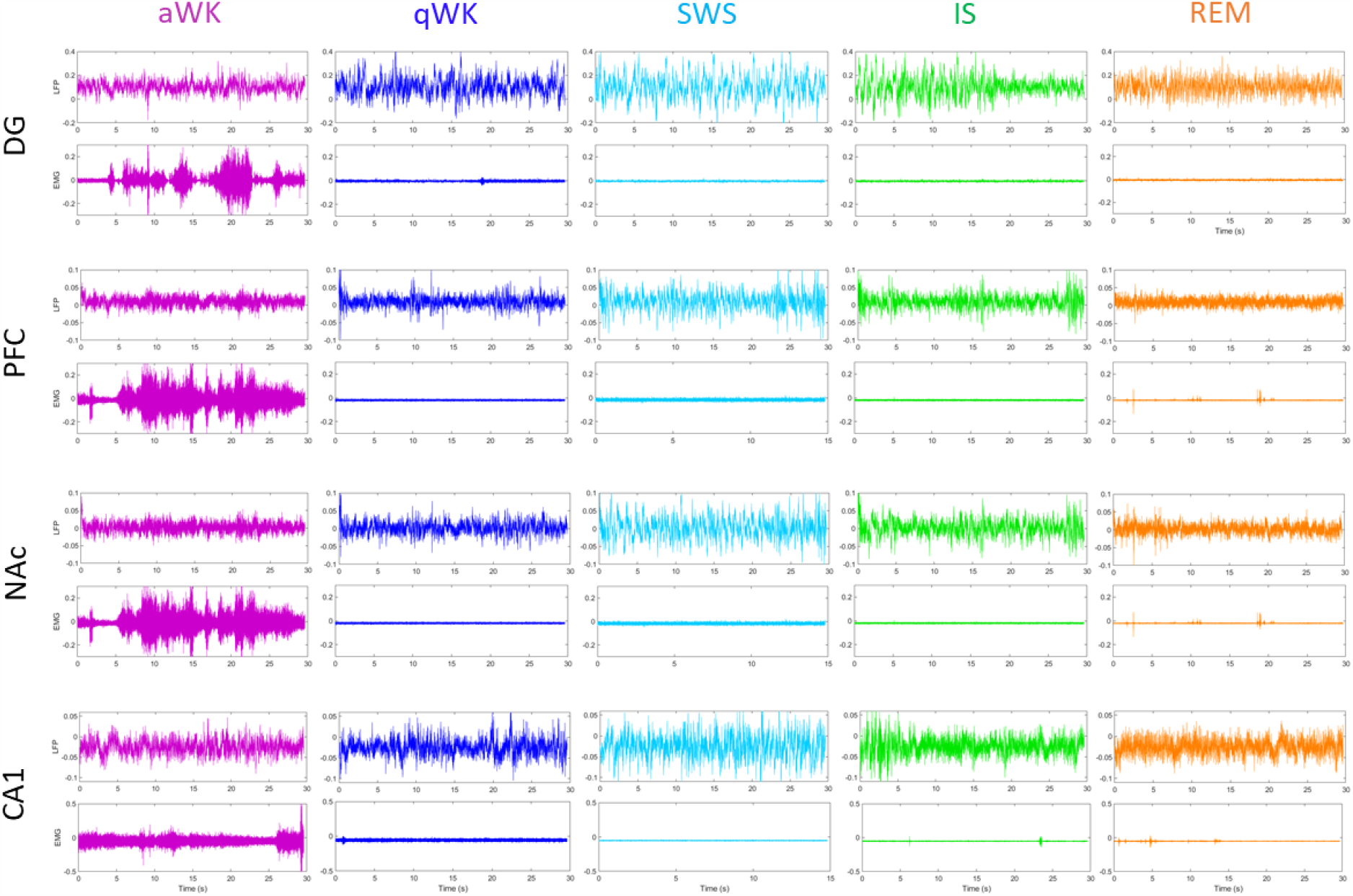
Vigilance states classification in different brain areas. Examples of LFP (top) and EMG (bottom) recordings for epochs of 30 s of active wakefulness (aWK), quiet wakefulness (qWK), slow wave sleep (SWS), intermediate sleep (IS) and rapid-eye-movement (REM) sleep in the Dentate Gyrus, prefrontal cortex (PFC), Nucleus Accumbens (NAc) or the CA1 region of the hippocampus (CA1).

**Figure S11:**
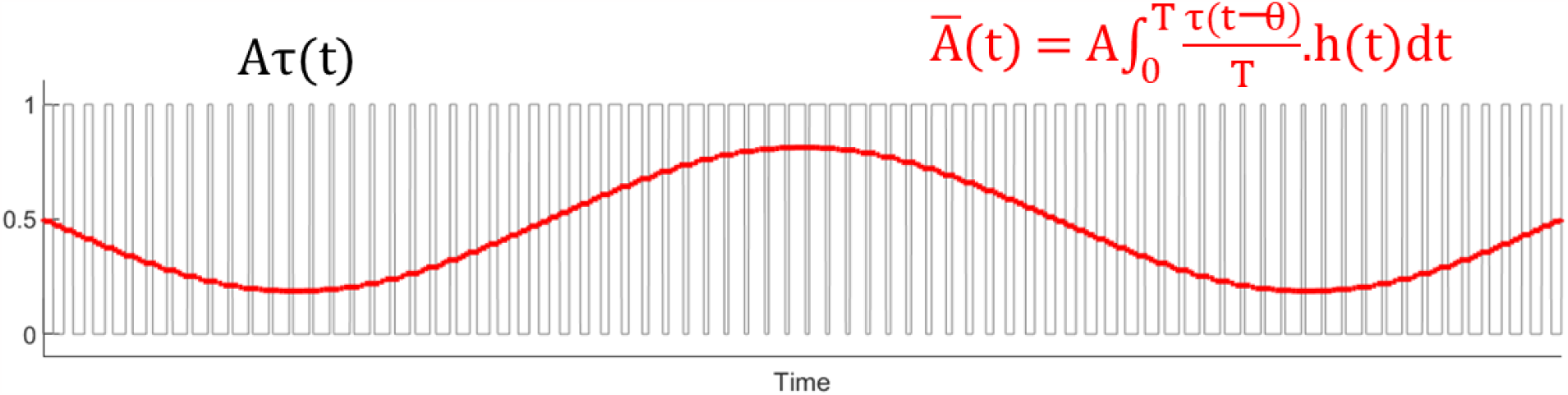
The average duration of a bistable state can encode slow variations, also called pulse width modulation. The black line represents a bistable signal with varying up or down states duration (analogy to the occurrence of a vigilance state). The red line represents the average amount of time spent in the up state computed over a sliding window.

**Figure S12:**
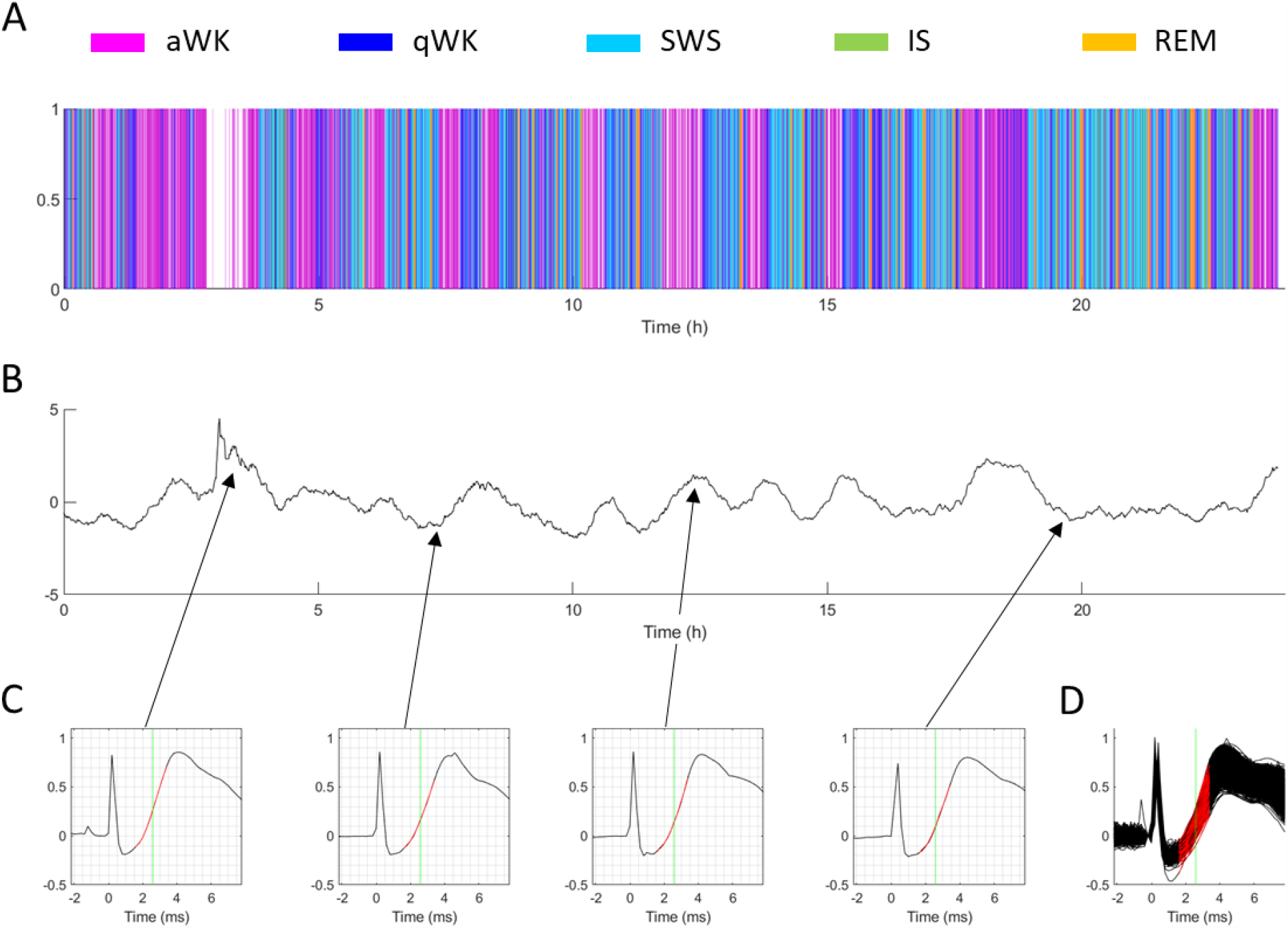
Representative examples of slope measurements at the PP-DG pathway of one animal across a day. (**A**) The black curves represent the normalized and detrended sliding average of synaptic response slopes for a representative animal. (**B**) Four examples of polynomial fits (red curve) of slopes on synaptic responses (black curve). (**C**) Stacking of all synaptic responses and their polynomial fits across 24h (n=2833).

**Figure S13:**
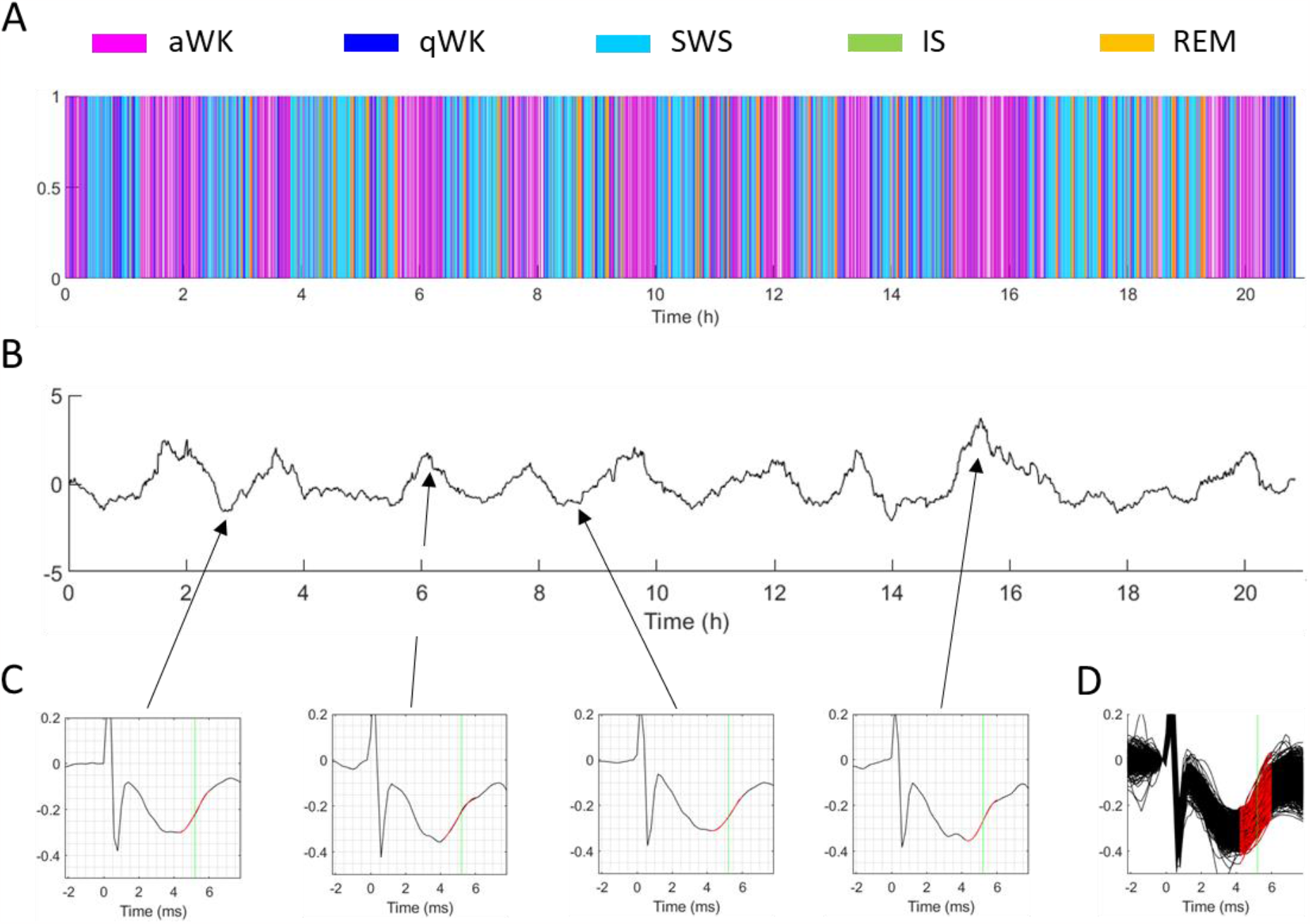
Representative example of slope measurements at the Fx-PFC pathway of one animal across a day. (**A**) The black curves represent the normalized and detrended sliding average of synaptic response slopes for a representative animal. (**B**) Four examples of polynomial fits (red curve) of slopes on synaptic responses (black curve). (**C**) Stacking of all synaptic responses and their polynomial fits across 21h (n=2502).

**Figure S14:**
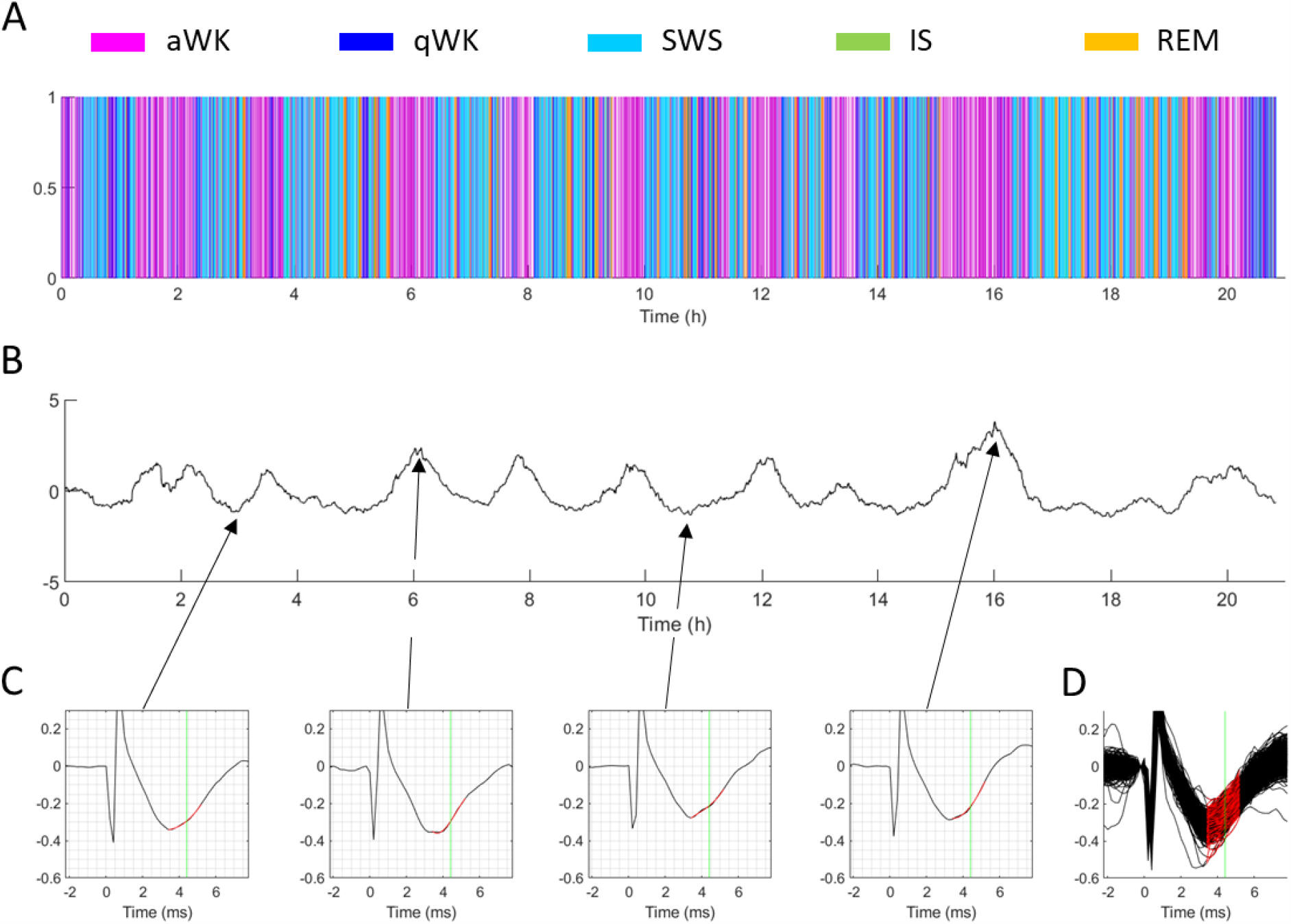
Representative example of slope measurements at the Fx-NAc pathway of one animal across a day. (**A**) The black curves represent the normalized and detrended sliding average of synaptic response slopes for a representative animal. (**B**) Four examples of polynomial fits (red curve) of slopes on synaptic responses (black curve). (**C**) Stacking of all synaptic responses and their polynomial fits across 21h (n=2171).

**Figure S15:**
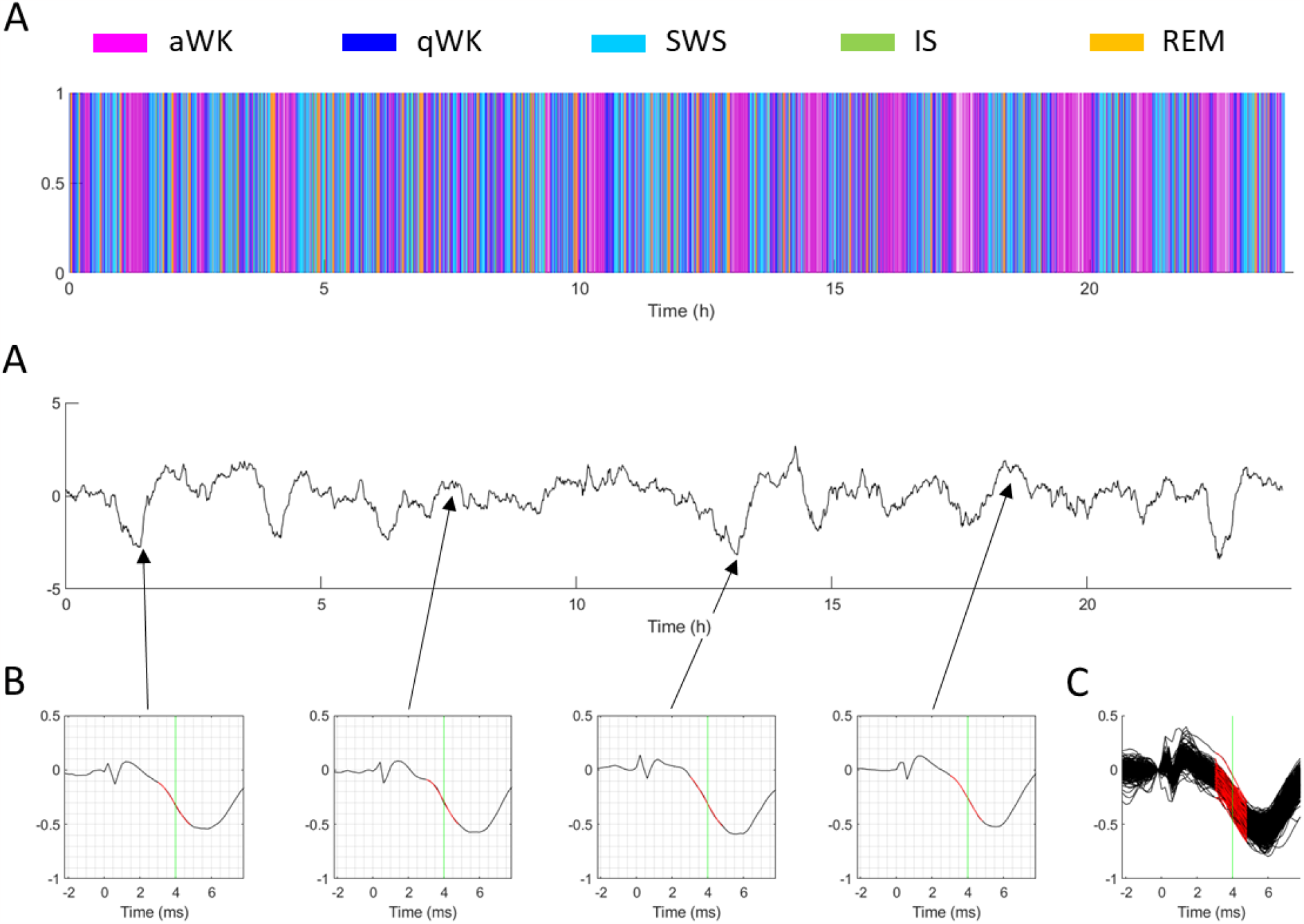
Representative example of slope measurements at the SC-CA1 pathway of one animal across a day. (**A**) The black curves represent the normalized and detrended sliding average of synaptic response slopes for a representative animal. (**B**) Four examples of polynomial fits (red curve) of slopes on synaptic responses (black curve). (**C**) Stacking of all synaptic responses and their polynomial fits across 24h (n=2814).

**Figure S16:**
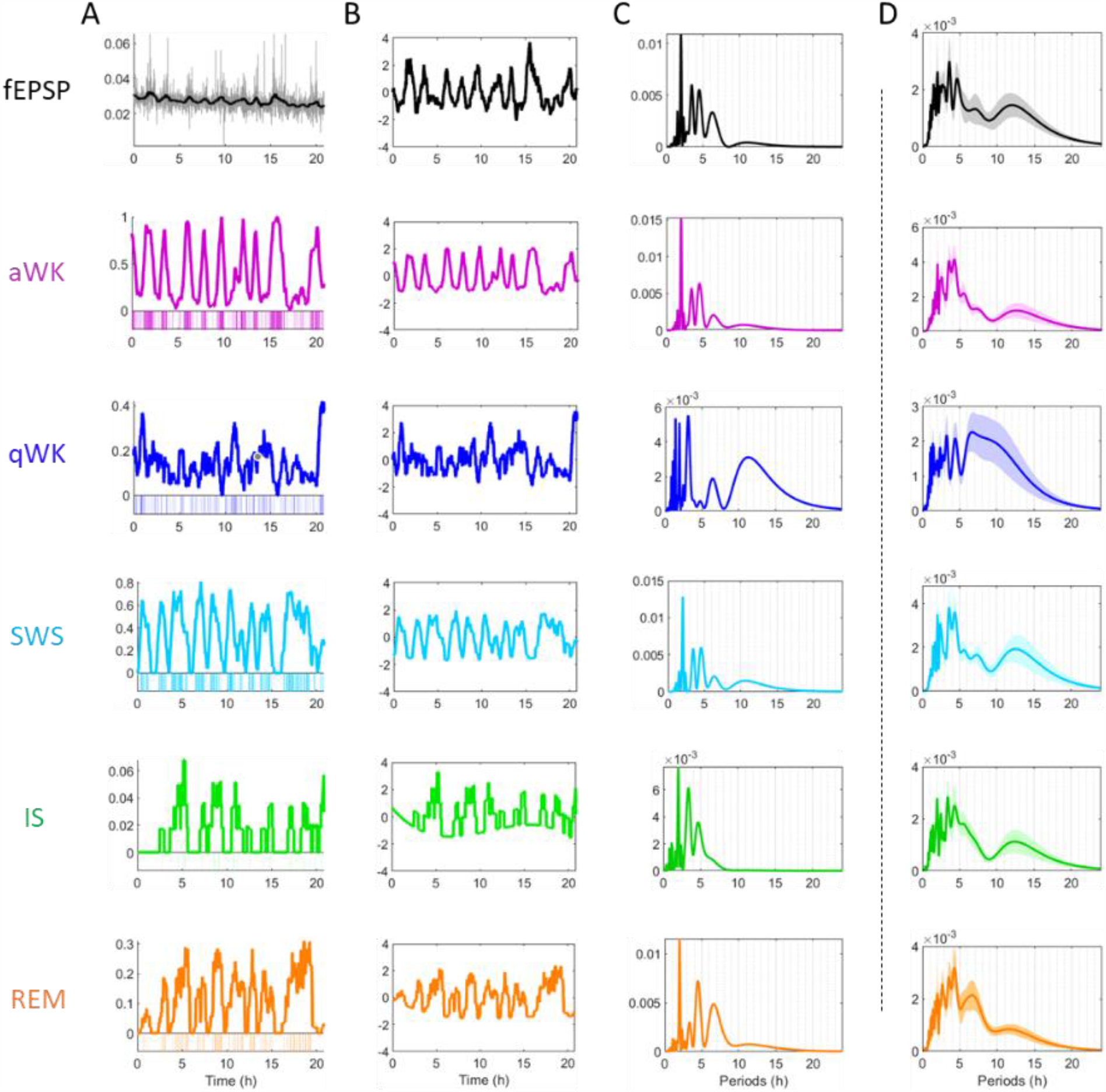
Synaptic response slopes at the Fx-PFC pathway and vigilance states share the same periodicities. (**A**), (**B**) From top to bottom: synaptic response slopes at the Fx-PFC pathway (fEPSP, in black) and vigilance states occurrence (aWK = active Wakefulness, in purple; qWK = Quiet Wakefulness, in dark blue; SWS = Slow Wave Sleep, in light blue; IS = Intermediate Sleep, in green; REM = Rapid Eye Movement Sleep, in orange) for a representative animal for 24 hours. (**A**) For synaptic slopes, the grey line represents raw measures, and the black line represents its sliding average over 1h. For vigilance states, the bottom histograms represent bouts of each state in color and missing values in grey. The colored lines represent the sliding averages of each vigilance state over 1h. (**B**) Black and colored lines represent normalized and detrended sliding averages. (**C**) Periodograms of the signals in B. (**D**) Mean periodograms for all rats (n=10). Shaded areas represent SEM.

**Figure S17:**
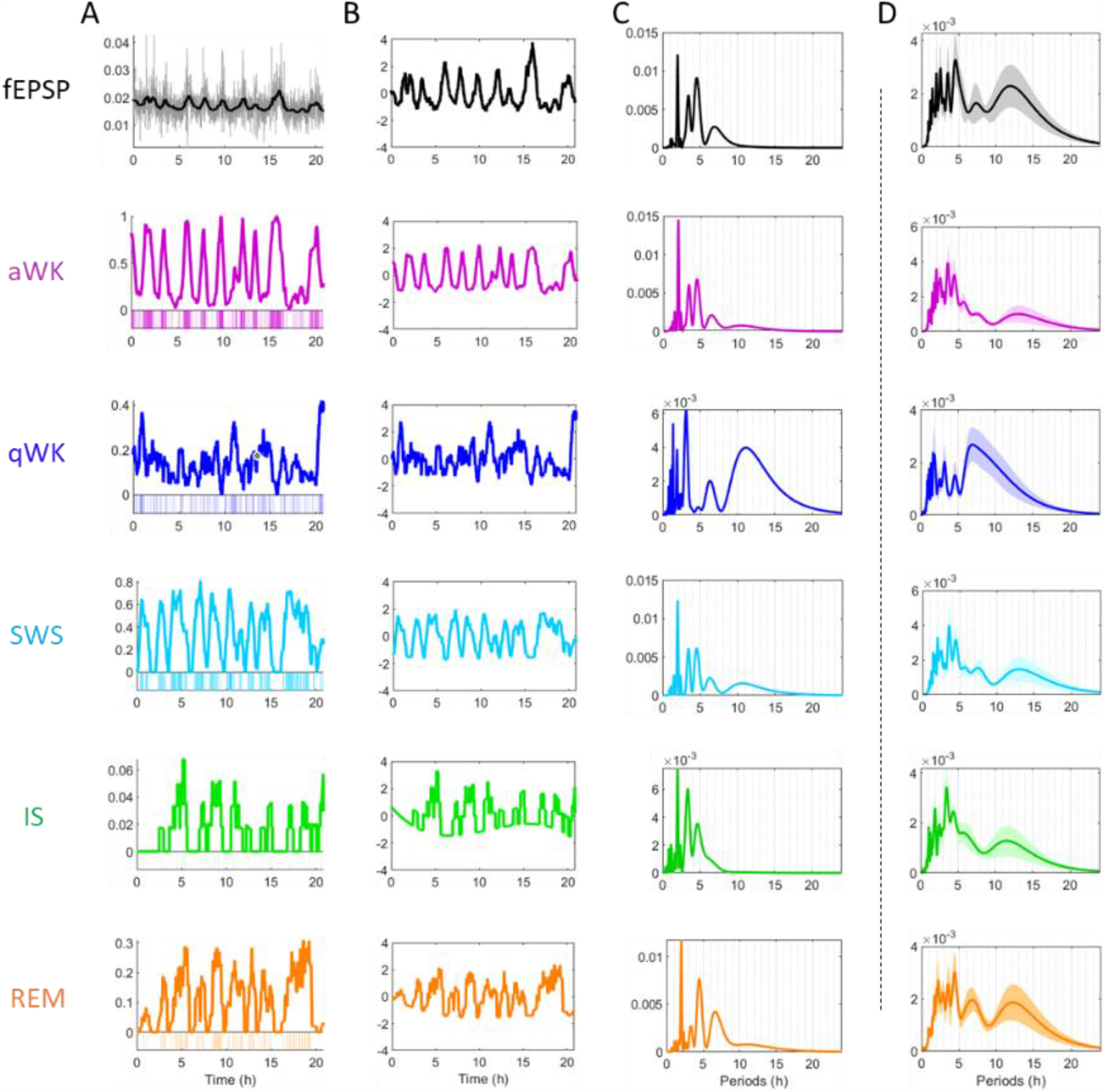
Synaptic response slopes at the Fx-NAc pathway and vigilance states share the same periodicities. (**A**), (**B**) From top to bottom: synaptic response slopes at the Fx-NAc pathway (fEPSP, in black) and vigilance states occurrence (aWK = active Wakefulness, in purple; qWK = Quiet Wakefulness, in dark blue; SWS = Slow Wave Sleep, in light blue; IS = Intermediate Sleep, in green; REM = Rapid Eye Movement Sleep, in orange) for a representative animal for 24 hours. (**A**) For synaptic slopes, the grey line represents raw measures, and the black line represents its sliding average over 1h. For vigilance states, the bottom histograms represent bouts of each state in color and missing values in grey. The colored lines represent the sliding averages of each vigilance state over 1h. (**B**) Black and colored lines represent normalized and detrended sliding averages. (**C**) Periodograms of the signals in B. (**D**) Mean periodograms for all rats (n=9). Shaded areas represent SEM.

**Figure S18:**
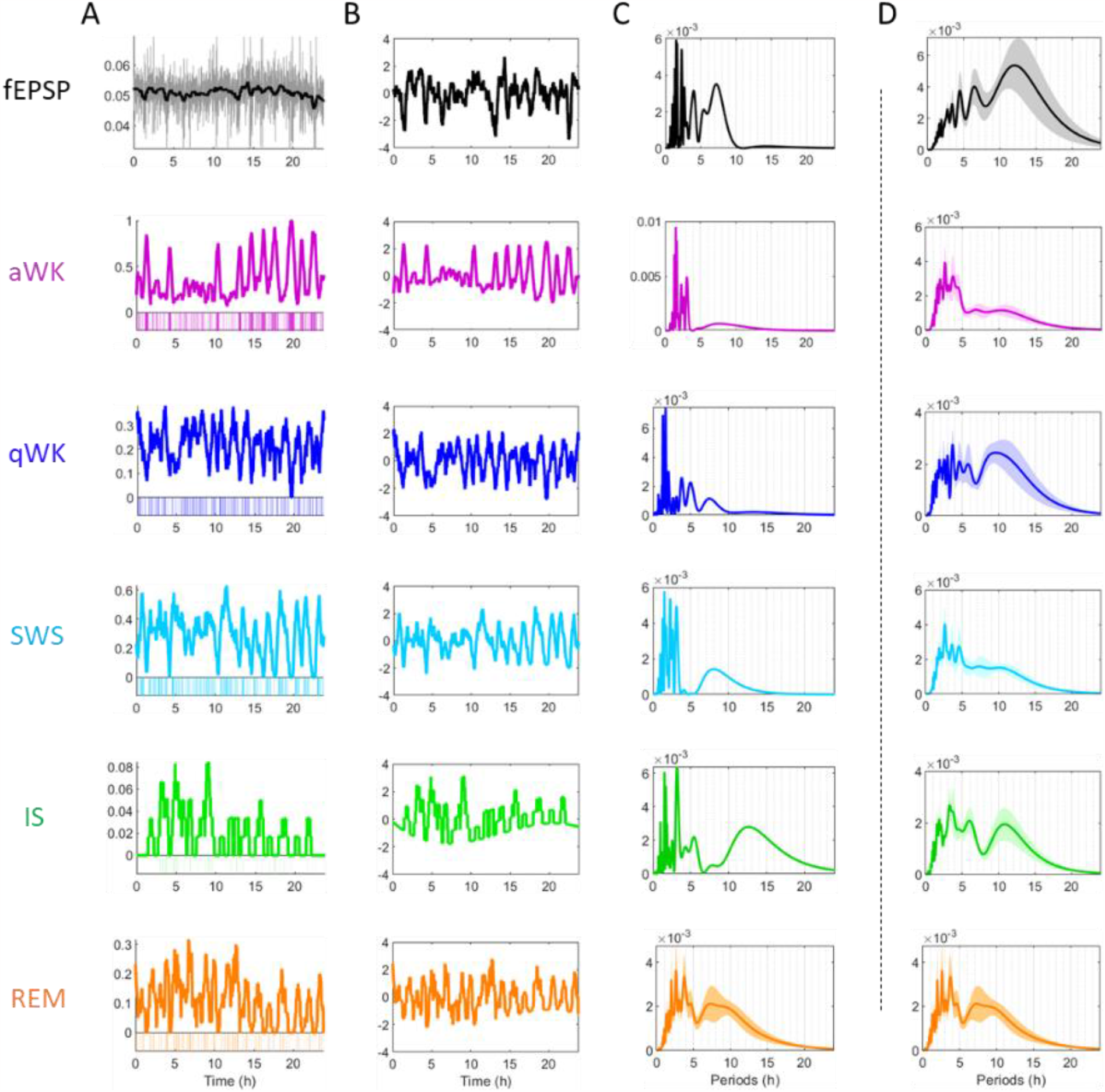
Synaptic response slopes at the SC-CA1 pathway and vigilance states share the same periodicities. (**A**), (**B**) From top to bottom: synaptic response slopes at the SC-CA1 pathway (fEPSP, in black) and vigilance states occurrence (aWK = active Wakefulness, in purple; qWK = Quiet Wakefulness, in dark blue; SWS = Slow Wave Sleep, in light blue; IS = Intermediate Sleep, in green; REM = Rapid Eye Movement Sleep, in orange) for a representative animal for 24 hours. (**A**) For synaptic slopes, the grey line represents raw measures, and the black line represents its sliding average over 1h. For vigilance states, the bottom histograms represent bouts of each state in color and missing values in grey. The colored lines represent the sliding averages of each vigilance state over 1h. (**B**) Black and colored lines represent normalized and detrended sliding averages. (**C**) Periodograms of the signals in B. (**D**) Mean periodograms for all rats (n=12). Shaded areas represent SEM.

**Figure S19:**
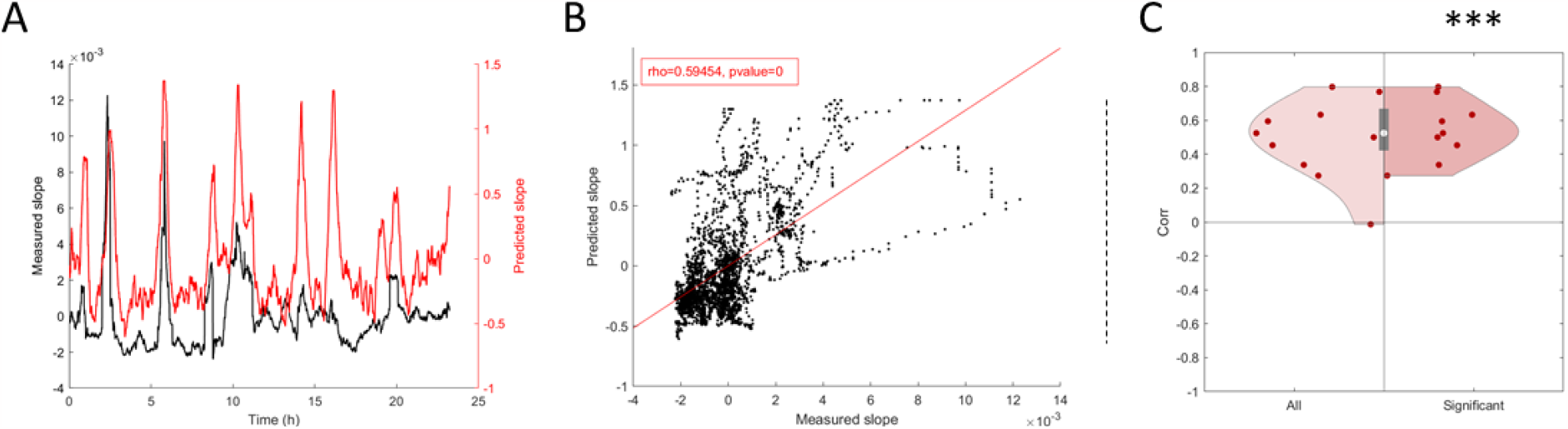
Vigilance state durations predict synaptic strength dynamics at the Fx-PFC pathway. See Figure 8 for legend (n=10).

**Figure S20:**
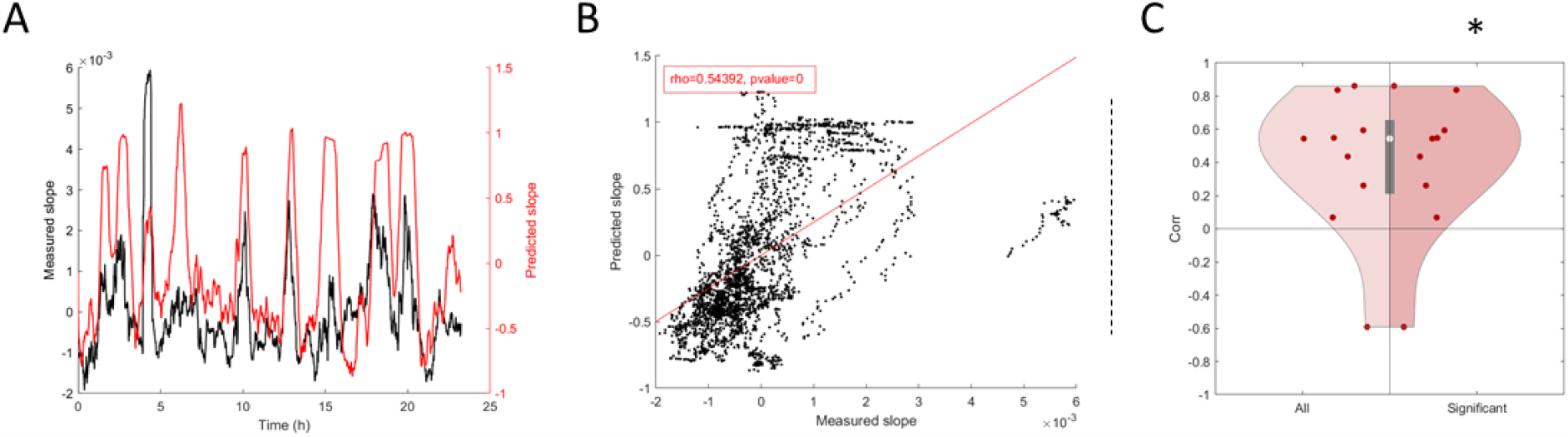
Vigilance state durations predict synaptic strength dynamics at the Fx-NAc pathway. See Figure 8 for legend (n=9).

**Figure S21:**
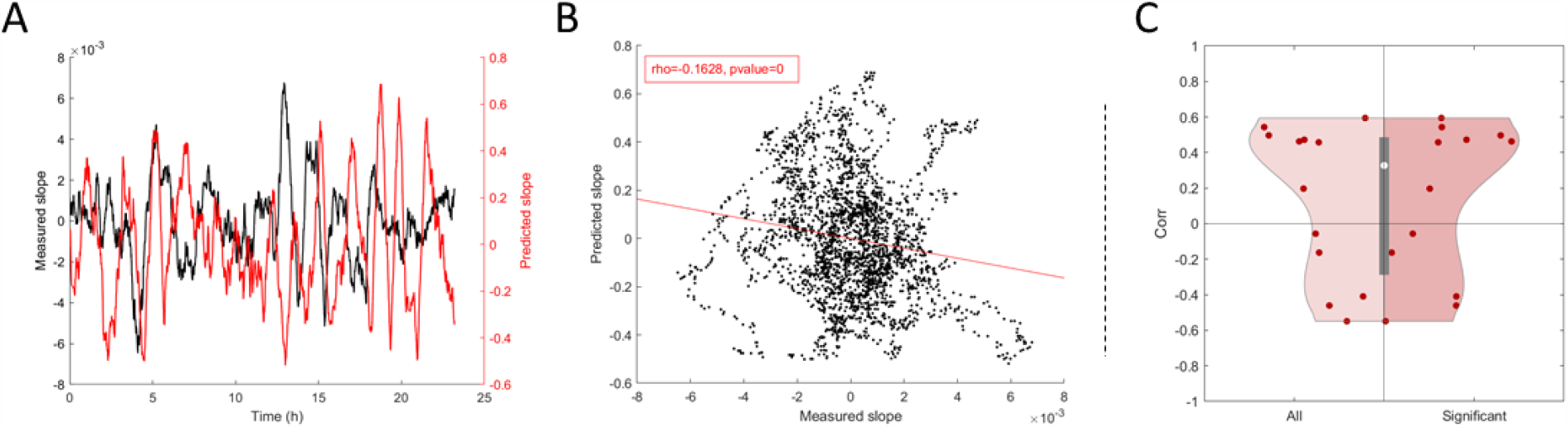
Vigilance states durations do not predict synaptic strength dynamics at the SC-CA1 pathway. See Figure 8 for legend (n=12).

**Figure S22:**
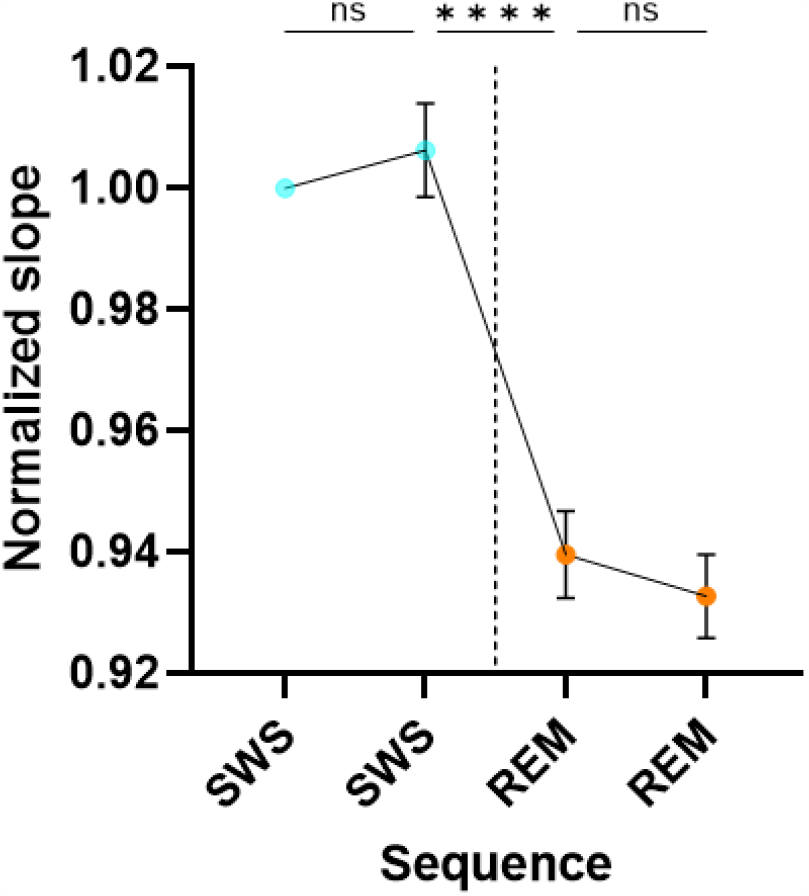
Synaptic response slopes variations in the DG at the transition between SWS and REM. To study the short-term effect of SWS to REM transition on synaptic strength, sequences of 2 consecutive SWS bouts followed by 2 consecutive REM bouts (n=550) were isolated in the hypnograms of rats implanted in the DG (n=35). For each sequence, synaptic response slopes were normalized to that of the first bout of the sequence. The graph represents the average normalized synaptic response slopes during SWS (in light blue) and during REM (in orange). Error bars represent SEM. Differences in synaptic response slopes were assessed by repeated measures 1 way ANOVA (p<0.001) and differences between consecutives bouts were assessed by Sidak’s multiple comparison’s test. ***: p<0.001.

**Figure S23:**
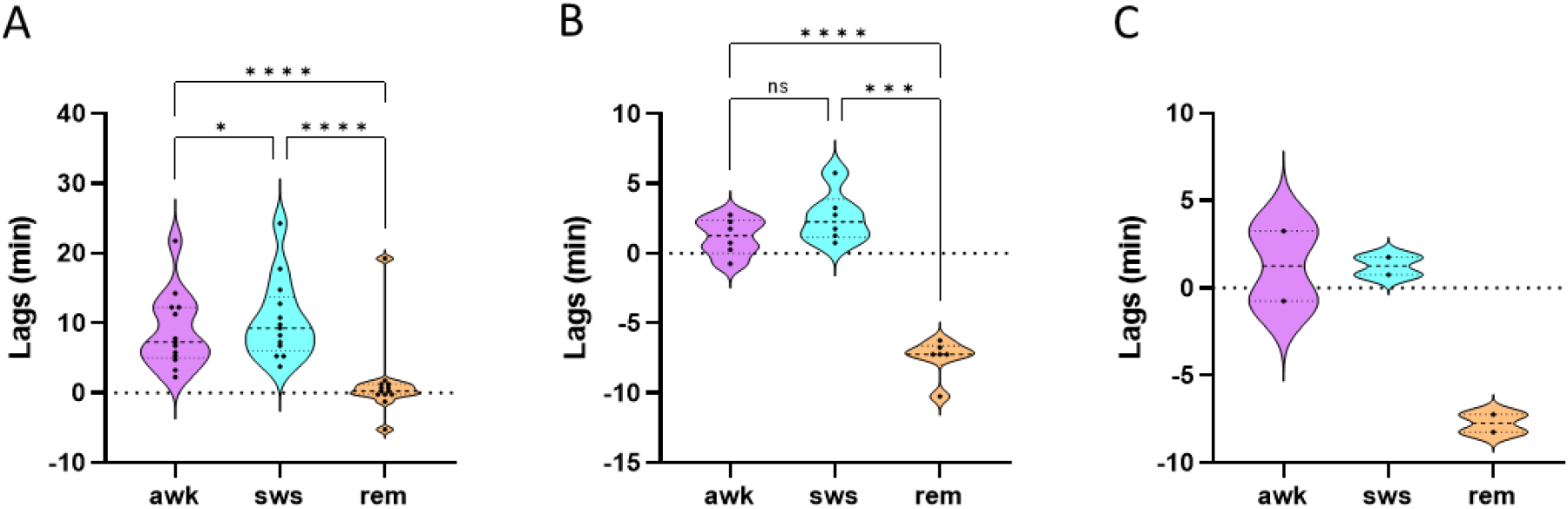
Slow Wave Sleep variations precede that of active Waking and REM sleep in their cross-correlations to synaptic response slopes. Comparisons of the time-lags in cross-correlation between each of the three main vigilance states (SWS, aWK and REM) and synaptic response slopes measured at (**A**) the PP-DG, (**B**) the Fx-PFC and (**C**) the Fx-NAc pathways. The comparison was performed on a subgroup of animals for which all the cross-correlation were significant to allow a paired comparison (n=14, n=6 and n=2, respectively).

**Table S1:** Statistics of cross-correlation coefficients at the PP-DG pathway.

| ***Vigilance States*** | ***Sample size*** | ***Significant sample size*** | ***Cross-correlation coefficients*** | ***p-value*** |
| --- | --- | --- | --- | --- |
| aWK | 35 | 18 | 0.76 | < 0.001 |
| qWK |  | 2 | -0.42 | NA |
| SWS |  | 17 | -0.69 | < 0.001 |
| IS |  | 20 | 0.51 | < 0.001 |
| REM |  | 22 | -0.66 | < 0.001 |

**Table S2:**
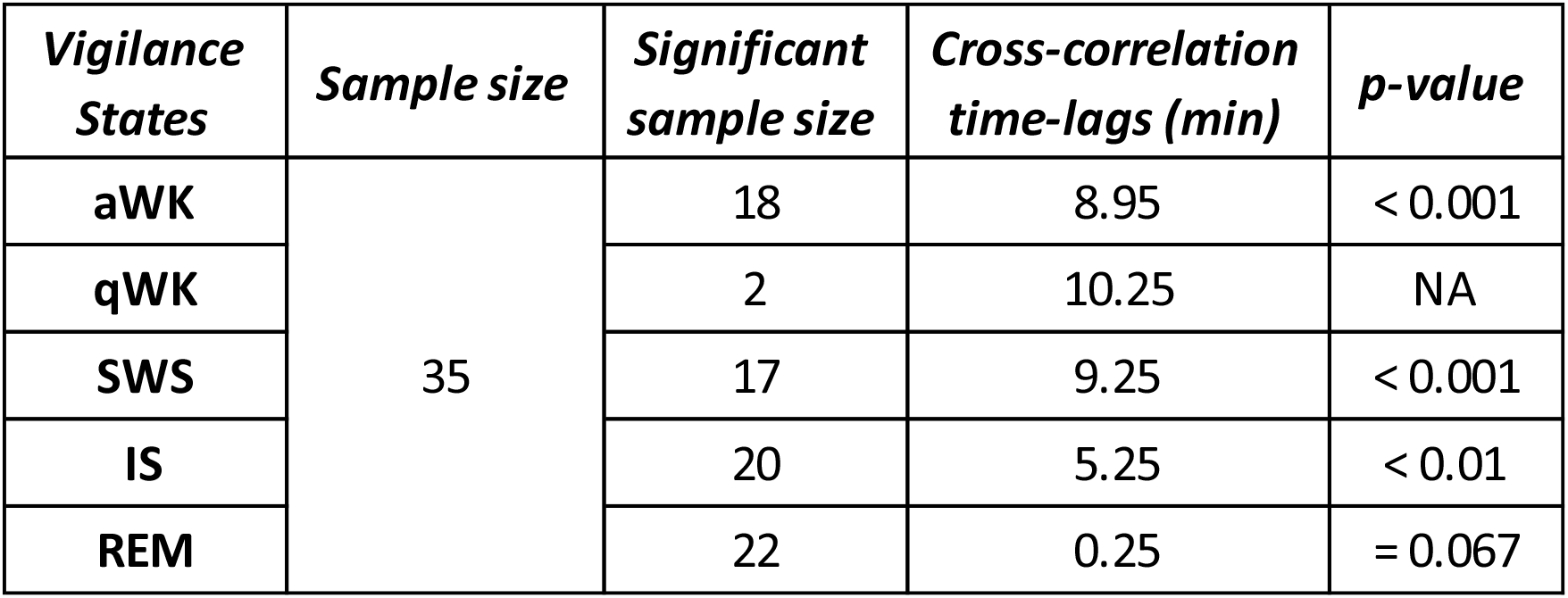
Statistics of cross-correlation time-lags at the PP-DG pathway.

**Table S3:** Statistics of cross-correlation coefficients at the Fx-PFC pathway.

| ***Vigilance States*** | ***Sample size*** | ***Significant sample size*** | ***Cross-correlation time-lags*** | ***p-value*** |
| --- | --- | --- | --- | --- |
| aWK | 10 | 8 | 0.66 | = 0.11 |
| qWK |  | 2 | 0.43 | NA |
| SWS |  | 8 | -0.65 | < 0.05 |
| IS |  | 8 | 0.33 | = 0.74 |
| REM |  | 6 | -0.50 | < 0.001 |

**Table S4:** Statistics of cross-correlation time-lags at the Fx-PFC pathway.

| ***Vigilance States*** | ***Sample size*** | ***Significant sample size*** | ***Cross-correlation time-lags (min)*** | ***p-value*** |
| --- | --- | --- | --- | --- |
| aWK | 10 | 8 | 1.25 | < 0.05 |
| qWK |  | 2 | -66.75 | NA |
| SWS |  | 8 | 2.25 | < 0.001 |
| IS |  | 8 | -61.15 | = 0.31 |
| REM |  | 6 | -7.25 | < 0.05 |

**Table S5:** Statistics of cross-correlation coefficients at the Fx-NAc pathway.

| ***Vigilance States*** | ***Sample size*** | ***Significant sample size*** | ***Cross-correlation coefficients*** | ***p-value*** |
| --- | --- | --- | --- | --- |
| aWK | 9 | 7 | 0.60 | = 0.16 |
| qWK |  | 0 | NA | NA |
| SWS |  | 5 | -0.67 | < 0.001 |
| IS |  | 5 | -0.12 | = 0.57 |
| REM |  | 5 | -0.54 | = 0.31 |

**Table S6:**
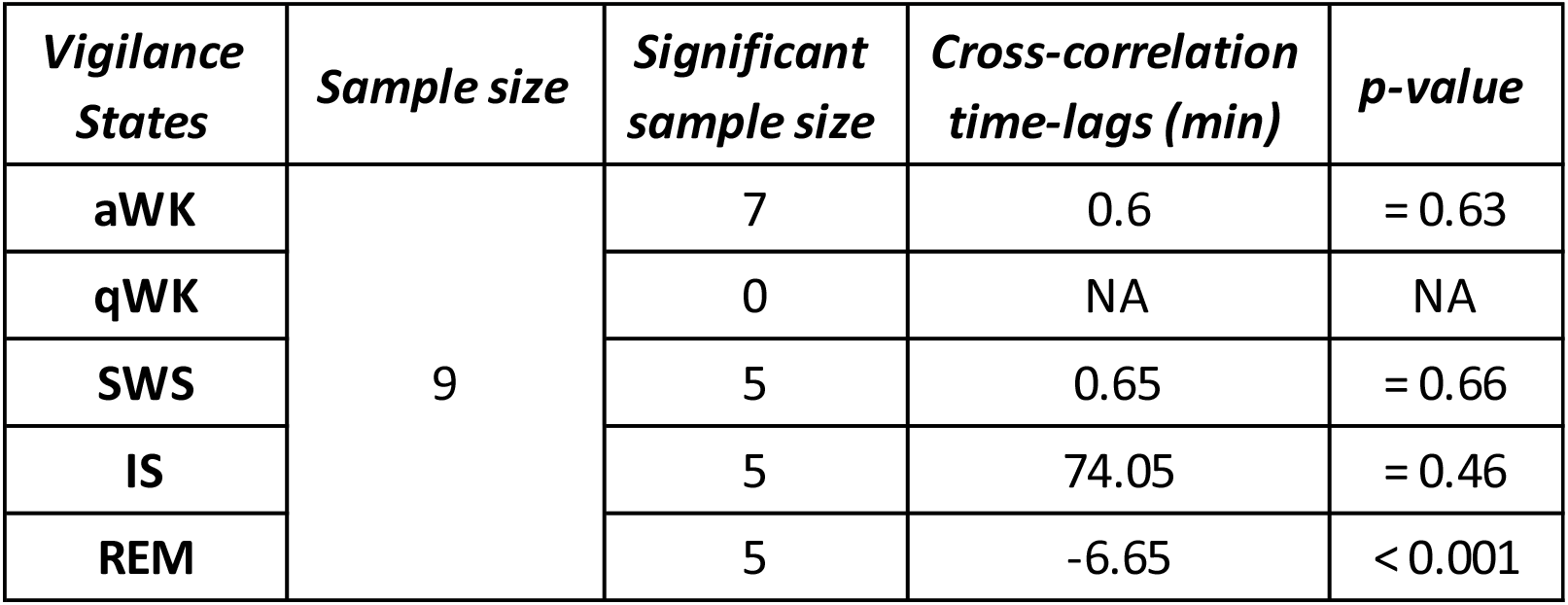
Statistics of cross-correlation time-lags at the Fx-NAc pathway.

**Table S7:** Statistics of cross-correlation coefficients at the SC-CA1 pathway.

| ***Vigilance States*** | ***Sample size*** | ***Significant sample size*** | ***Cross-correlation coefficients*** | ***p-value*** |
| --- | --- | --- | --- | --- |
| **aWK** | 12 | 3 | -0.24 | = 0.63 |
| **qWK** |  | 2 | 0.064 | NA |
| **SWS** |  | 4 | 0.64 | = 0.38 |
| **IS** |  | 1 | -0.36 | NA |
| **REM** |  | 1 | -0.53 | NA |

**Table S8:** Statistics of cross-correlation time-lags at the SC-CA1 pathway.

| ***Vigilance States*** | ***Sample size*** | ***Significant sample size*** | ***Cross-correlation time-lags (min)*** | ***p-value*** |
| --- | --- | --- | --- | --- |
| **aWK** | 12 | 3 | 0.59 | = 0.77 |
| **qWK** |  | 2 | 22.25 | NA |
| **SWS** |  | 4 | 1.63 | = 0.30 |
| **IS** |  | 1 | 460.25 | NA |
| **REM** |  | 1 | 17.25 | NA |

**Figure S24:**
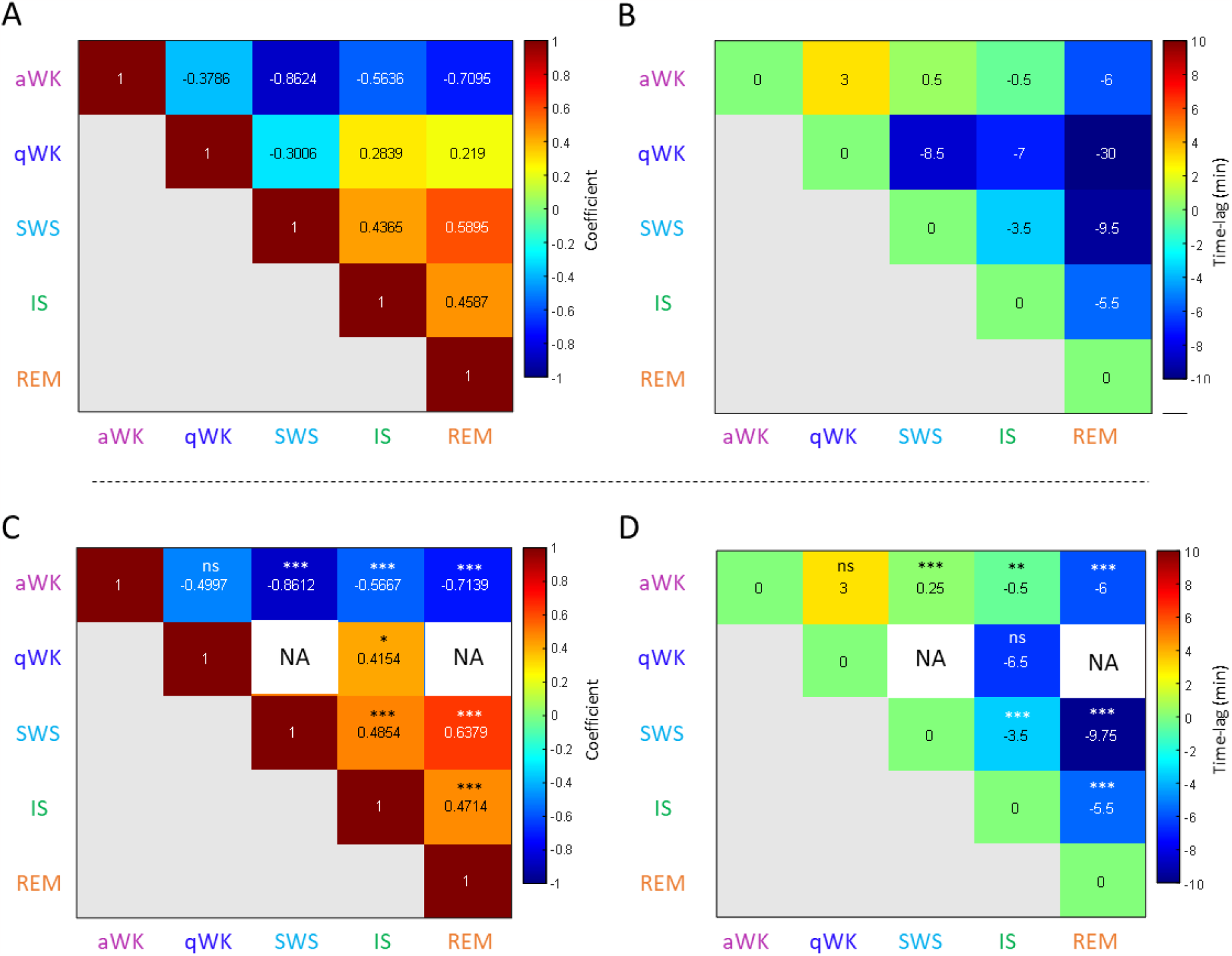
Cross-correlations between vigilance states. (**A**,**C**) Matrices of median cross-correlations coefficients between pairs of vigilance states of animals implanted in the Dentate Gyrus (n=35). (**B**,**D**) Matrices of median time-lags in the cross correlations of animals implanted in the Dentate Gyrus (n=35). (**A**-**B**) Cross correlation coefficients and time-lags for all animals. (**C**-**D**) Non-spurious only (>CCSL) cross correlation coefficients and corresponding time-lags. For example, we show that REM sleep is delayed by a median of 9.75 min compared to SWS. Statistical significance difference to a theoretical median of 0 is represented. *: p<0.05, **: p<0.01, ***: p<0.001, ns: non-significant. NA: too few cross-correlation coefficients passed the CCSL to perform statistical analysis.

## Notes

### Competing Interest Statement

The authors have declared no competing interest.

